# Engineering T cell resistance to HIV-1 infection via knock-in of peptides from the heptad repeat 2 domain of gp41

**DOI:** 10.1101/2021.04.30.442168

**Authors:** Alexandra Maslennikova, Natalia Kruglova, Svetlana Kalinichenko, Dmitriy Komkov, Mikhail Shepelev, Dmitriy Golubev, Andrei Siniavin, Andrei Vzorov, Alexander Filatov, Dmitriy Mazurov

## Abstract

Previous studies suggest that short peptides from the heptad repeat 2 (HR2) domain of gp41 expressed on the cell surface are more potent inhibitors of HIV-1 entry than soluble analogs. However, their therapeutic potential has only been examined using lentiviral vectors. Here, we aimed to develop CRISPR/Cas9-based fusion inhibitory peptide knock-in (KI) technology for the generation and selection of HIV-1-resistant T cells. First, we cloned a series of HIV-1 fusion inhibitory peptides and embedded them in CD52, the shortest GPI-anchored protein, which efficiently delivers epitope tags to the cell surface and maintains a sufficient level of KI. Among the seven peptides tested, MT-C34, HP-23L, and 2P23 exhibited significant activity against both cell-free and cell-to-cell HIV-1 infection. Unlike membrane-bound peptides, the shed variant of MT-C34 provided insufficient protection against HIV-1 due to its low concentrations in the culture medium. Using Cas9 plasmids or ribonucleoprotein electroporation and cell sorting with antibodies raised against gp41 peptides, we generated CEM/R5 cells with biallelic KI of MT-C34 (embedded in CD52 for expression in lipid rafts) and 2P23 (N-terminally fused to CXCR4). In combination, these peptides provided a higher level of protection than individual KI. By extending homology arms and substituting PCR donor DNA with a plasmid containing signals for nuclear localization, we achieved KI of MT-C34 into *CXCR4* loci and HIV-1 proviral DNA at levels of up to 35% in CEM/R5 cells and increased KI occurrence from undetectable to 4-5% in CD4 lymphocytes. Comparative analysis of lentiviral and HDR-based delivery strategies showed that KI led to the higher MT-C34 expression and stronger protection of primary CD4 lymphocytes from HIV-1 than lentiviral transduction, albeit the efficiency of KI needs further improvements in order to meet clinical requirements. Thus, the developed CRISPR/Cas9 platform offers a new opportunity for antiviral peptide delivery with a concomitant precise genetic modification of targeted locus that can be employed to strengthen cell protection against HIV.

**AUTHOR SUMMARY:** HIV is a human lentivirus that infects CD4-positive immune cells and, when left untreated, manifests in the fatal disease known as acquired immunodeficiency syndrome. Antiretroviral therapy (ART) is not leading to viral clearance, and HIV persists in the organism as a latent provirus. One way to control infection is to increase the population of HIV-resistant CD4 lymphocytes via entry molecule knockout or expression of different antiviral genes. Peptides from the heptad repeat (HR) domain of gp41 are potent inhibitors of HIV-1 fusion, especially when designed to express on the cell surface. Individual gp41 peptides encoded by therapeutic lentiviral vectors have been evaluated and some have entered clinical trials. However, a CRISPR/Cas9-based gp41 peptide delivery platform that operates through concomitant target gene modification has not yet been developed due to low knock-in (KI) rates in primary cells. Here, we systematically evaluated the antiviral activity of different HR2-peptides cloned into the shortest carrier molecule, CD52. The resulting small-size transgene constructs encoding selected peptides, in combination with improvements to enhance donor vector nuclear import, helped to overcome precise editing restrictions in CD4 lymphocytes. Using KI into CXCR4, we demonstrated different options for target gene modification, effectively protecting edited cells against HIV-1.

## INTRODUCTION

Human immunodeficiency virus type I (HIV-1) is a retrovirus, the genome of which stably integrates into the host’s DNA during viral infection. As such, HIV-1 eradication is only possible when all infected cells in the host organism are either killed or cured via “DNA surgery.” The first instance of HIV being cured was reported after the allogeneic transplantation of CCR5(Δ32) bone marrow stem cells to a patient in Berlin (1, 2). Later on, a remission after analytical treatment interruption for at least 30 months was observed in the London patient after a similar transplantation (3). In other cases, however, engraftments were not as successful, and patients experienced viral rebounds (4). The Berlin patient’s case inspired researchers to develop genetic tools to make cells resistant to HIV. Zinc finger nucleases (ZFNs) specific to CCR5 were the first programmed genomic nucleases to enter HIV clinical trials (5). With the adaptation of a bacterial defense system CRISPR-Cas9 (clustered regularly interspaced short palindromic repeats and CRISPR-associated protein 9) for gene editing in mammalian cells (6, 7), diverse genetic manipulations, including HIV proviral excision (8, 9) and/or inactivation (10, 11) and CCR5 and CXCR4 knockout (12-14), have been found to be effective and easily achievable. However, three major problems related to CRISPR-Cas9-based editing—emergence of viable HIV escape mutants during indel formation (10, 15, 16), off-target activity of Cas9 (17, 18), and low efficiency of precise genome modification via homology-directed repair (HDR) (19, 20)—create a serious barrier to the use of CRISPR in clinical practice.

Of the various anti-HIV genetic strategies, the inhibition of viral entry is one of the most efficient. Short peptides from the heptad repeat 2 (HR2) domain of gp41 inhibit HIV entry by interacting with the HR1 region of gp41, blocking the formation of gp41 6-helix bundles required for membrane fusion (21). The first generation of HR2 peptides consisted of relatively long T-20, C36, and C46 peptides to which the virus rapidly evolved resistance. Subsequent modifications aimed to 1) enhance the interaction of the peptide with the HR1 deep hydrophobic pocket by adding an M-T hook (22, 23) or introducing mutations that stabilized the α-helices (24, 25), or 2) broaden protection against HIV-2 and SIV (26), resulting in the development of a second and third generations of short peptides with higher inhibitory potency and lower viral resistance. To date, only two fusion inhibitor peptides have been approved for clinical use: enfuvirtide (T-20), approved by the FDA, and sifuvirtide, approved for use in China. While synthetic gp41 peptides and their chemical modifications have been studied extensively, much less is known about the potency of genetically encoded analogs. The first studies from Dorothee von Laer’s lab showed that T-20 and C46 peptides expressed on the cell surface in the context of the human low-affinity nerve growth factor receptor (LNGFR) are very potent inhibitors of HIV-1 entry (27, 28). In 2016, Liu et al. reported that a shorter C34 peptide linked to a GPI-attachment signal from the decay-accelerating factor (DAF) and expressed in lipid rafts confers strong cellular resistance to HIV-1 infection (29). Recently, He’s lab demonstrated that the third-generation peptide 2P23, when expressed in a similar way, fully protects cells against HIV-1, HIV-2, and SIV infection (30). Besides transmembrane linking or GPI-anchoring, direct fusion of C34 to the N-termini of CXCR4 and CCR5 has been reported to confer cells with a strong resistance to HIV infection *in vitro* and *in vivo* (31). Collectively, these data indicate that gp41 peptides, when expressed on a plasma membrane and localized in lipid rafts or co-receptor sites where HIV fusion occurs, become very active in comparison to the soluble form. However, in all these studies, peptides were expressed from retro- or lentiviral vectors with no aim of adapting them for knock-in into endogenous genes. Since HDR-based gene editing is sensitive to the size of the DNA insert, the shortest construct comprised of a peptide and a carrier molecule is desirable in order to achieve a reasonable level of precision in the genetic modification.

We have previously described a novel method for gene-edited cell selection called SORTS (surface oligonucleotide knock-in for rapid target selection), which is based on the targeted knock-in of epitope tags embedded in CD52 (32). CD52 is the shortest human GPI-anchored protein that can effectively deliver peptides to the plasma membrane, where they serve as makers for fluorescence-activated cell sorting (FACS) isolation of edited cells. In the present study, we modified SORTS by substituting the epitope tag in CD52 with a peptide from the HR2 domain of gp41. We selected the shortest and most potent soluble peptides of the last generation described to date and engineered lentiviral vectors to stably express these peptides on the cell surface. Of the seven tested peptides, MT-C34, HP23L, and 2P23 demonstrated the highest activity against both cell-free and cell-to-cell infection by HIV-1. In contrast, modifications that aimed to secrete, shed, or noncovalently link the MT-C34 peptide to HIV chemokine co-receptors did not improve modified or bystander cell protection. Using CRISPR-Cas9 co-knock-in of MT-C34 and 2P23 (or HP23L) into the endogenous *CXCR4* gene followed by the sorting of double-positive cells labeled with anti-MT-C34 and anti-2P23(HP23L) antibodies (Abs), we isolated a pure population of CEM CD4^+^ T cells, where the indicated peptides were N-terminally fused to CXCR4 or expressed separately in a GPI-linked form. Both genetic modifications preserved CXCR4 function and completely protected cells from different clades of HIV-1 with various tropisms, as well as from gp41 mutant viruses. Through the optimization of donor DNA, i.e., the extension of homology arms and substitution of the PCR donor with a plasmid, we achieved ∼30% HDR efficiency at the *CXCR4* locus in CEM cells. We also achieved a reasonable level of peptide KI into human primary cells using modifications aimed to enhance the nuclear import of donor DNA and demonstrated that MT-C34 KI resulted in stronger protection of CD4 lymphocytes in comparison to lentiviral transduction. Thus, unlike vector-based HIV therapies, this approach relies on precise genetic modification with CRISPR-Cas9 that enables the expression of protective peptides, modification of cellular receptors and isolation of modified cells.

## RESULTS

### Construction and antiviral activity of CD52-embedded gp41 peptides

We have previously demonstrated that SORTS can be adapted for the isolation of T cell population with precisely inactivated HIV-1 genome via the KI of a CD5HA2 marker construct, in which the peptide sequence SQTSSP, located in the middle of CD52, is replaced with an HA tag (32). We hypothesized that providing the selected cells with a protective peptide would render “cured” cells resistant to repeat infection with HIV. We selected several peptides from HR2 of gp41 with high and broad anti-HIV activity as reported in the literature and substituted the HA sequence in CD5HA2 constructs with one of the peptides specified in Figure 1A. We then jointed it to the upstream green fluorescent protein mClover via a P2A sequence for separate translation. Three peptides—T-20, C34, and C24 (an inactive form used as a negative control that represents the overlapped sequence of the first two peptides without a pocket-binding domain (33))—were the natural sequences from the HxB2 strain of HIV-1. The other five peptides had the following modifications: MT-C34 had M-T hook residues (23, 34) while P52 (35), MT-WQ-IDL (36), HP23L (37, 38), and 2P23 (26, 30) contained α-helix stabilizing mutations with or without an additional hook-like structure at the N- or C-terminus. To test the inhibitory activity of the peptides, the 293T/CD4/R5 cells—permissive to CCR5-and CXCR4-tropic HIV—were first transduced with one of the generated lentiviral constructs at MOI <0.3 and sorted based on mClover fluorescence. As the T-20 peptide was not properly expressed in our system (likely due to its high C-terminal hydrophobicity), it was excluded from all subsequent experiments. 293T cells with or without receptor/co-receptor/peptide expression were infected with an equal quantity of viral particles pseudotyped with R5-tropic JRFL, ZM135, or X4-tropic NL4-3 Env (referred to below as pseudoviruses (PVs)). Levels of HIV-1 infection were quantified using previously developed replication-dependent inLuc vector (39, 40). The luciferase gene in this vector is interrupted by an intron and together with CMV promoter and polyA signal placed in the reverse orientation relative to the viral genomic sequence. In producer (transfected) cells, U3-driven reporter RNA is spliced and packaged. Followed by infection of target cells and reverse transcription, the expression cassette becomes activated, and after integration in the host genome, the functional reporter protein can be expressed. The in-Luc reporter was developed to measure HIV-1 infectivity in mixed producer–target cell cultures but can be concurrently used for cell-free infectivity measurement. As shown in Fig 1B, P52 and MT-WQ-IDL demonstrated less potent inhibition (60–79% and 79–93% depending on Env, respectively) than other peptides, which blocked HIV-1 cell-free infection at levels of 98% or higher. The most protective peptides—C34, MT-C34, HP23L, and 2P23—were further tested in cell-to-cell infection. For this purpose, we transduced Raji/CD4/R5 cells with lentiviruses expressing these peptides and sorted them as outlined above. We then measured the levels of infection from non-permissive Raji producer cells to the Raji/CD4/R5 cells with or without peptides through inLuc transduction, as described above (40). Similar to cell-free infection, C34 and MT-C34 also actively inhibited HIV-1 cell-to-cell infection, while HP23L and 2P23 displayed less potent inhibition (Fig. 1C). Ultimately, we selected four peptides—C34, MT-C34, HP23L, and 2P23, embedded in a CD52 carrier molecule—with the highest potencies to inhibit HIV-1 entry.

**Figure 1.**
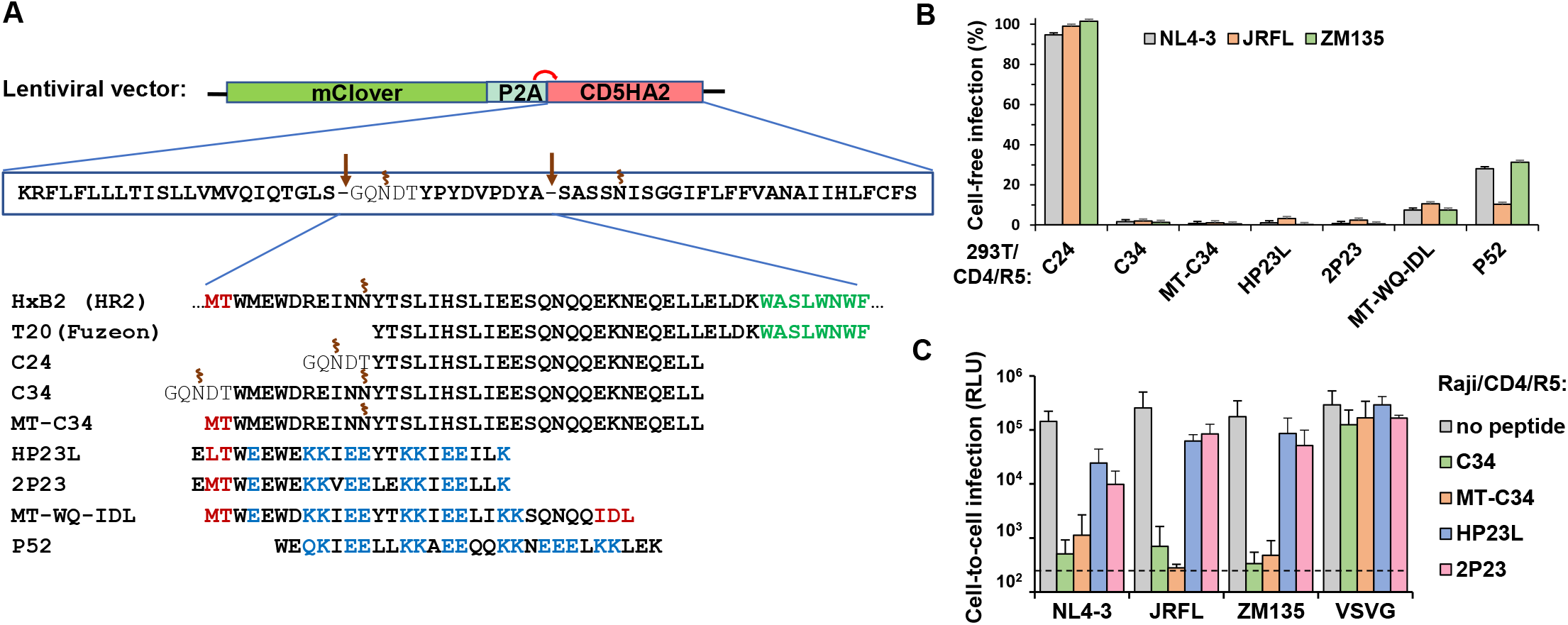
Construction and anti-HIV-1 activity of peptides from HR2-domain of gp41 expressed in the context of CD52. **A**. Schematic of generated lentiviral constructs for stable expression of gp41 peptides. Amino acids forming hook-like structure, α-helix-stabilizing mutations, and hydrophobic parts of peptides are highlighted in red, blue and green, respectively. Sites of proteolytic cleavage and N-glycosylation are indicated by arrows and spirals, respectively. **B**. Effects of GPI-anchored peptides on HIV-1 cell-free infection. 293T/CD4/R5 cells with the stable expression of indicated peptides were infected by the equal doses of HIV-1 VLPs pseudotyped with Envs from different strains of HIV-1 or VSV. Forty eight hours later, the levels of inLuc reporter transduction were measured. The infectivity values obtained for HIV-1 Envs were first adjusted to the respective infectivity data detected with VSV-G; the resulting value estimated in 293T/CD4/R5 cells without peptides was set at 100%, and the levels of infection for cells carrying indicated peptides were recalculated relative to that value. **C**. Effects of selected peptides on cell coculture infection. Raji cells were transfected to produce VLPs pseudotyped with indicated Envs and then directly mixed to the indicated Raji/CD4/R5 target cells at 1:1 ratio. The levels of infectivity were estimated by measuring inLuc transduction in 54 h of coculture and presented as raw data. The background level of Luc activity is shown by dashed line. The data in B and C are representative of at least three independent measurements and shown as average values ± standard deviations.

### Analysis of gp41 peptides and HIV receptor/co-receptor expression levels on the surfaces of transduced cells

In order to verify whether the inhibition of HIV-1 entry depends on the level of peptide and/or viral receptor/co-receptor expression on target cells, we generated mouse monoclonal antibodies (mAbs) against C24 (which could also recognize C34 and MT-C34 peptides) and HP23L, as well as a rabbit polyclonal Ab specific to the 2P23 peptide. 293T/CD4/R5 and Raji/CD4/R5 cells with or without peptides were individually labeled with the Abs indicated above or mAbs specific to CD4, CXCR4, and CCR5 and analyzed by flow cytometry. As expected, MT-WQ-IDL, P52, HP23L, and 2P23 were not recognized by anti-C24 mAbs, whereas the peptides C24, C34, and MT-C34 were intensely stained on the surface of 293T cells (Fig. 2A and B). Similarly, the anti-HP23L mAb and anti-2P23 Ab demonstrated specificity in peptide recognition. Unlike C34-based peptides, which had high levels of expression on both 293T and Raji cells, the HP23L and 2P23 peptides were expressed less intensely, especially on Raji cells (Fig. 2C). These two peptides lack additional sites of N-glycan attachment, which are present in C24, C34, and MT-C34 sequences (Fig. 1A). This suggests that the N-glycosylation of chimeric CD52 proteins at multiple sites may stimulate the export of carried peptides to the plasma membrane, which has been shown in our previous work (32) and similar studies (41, 42). The non-optimal expression of HP23L and 2P23 embedded in CD52 may lower their antiviral potency, as we observed in a cell co-culture infectivity test (Fig. 1C); however, the type of peptide can also determine its activity. Monitoring HIV-1 receptor and co-receptor expression levels on peptide transduced cells did not reveal any substantial differences between them, suggesting that the inhibitory activity of the peptides was not mediated by the downregulation of viral entry molecules. All HIV-1 entry molecules were expressed on transduced cells at comparably high levels, except for CXCR4 on 293T cells, the endogenous level of expression of which is known to be low but sufficient for HIV-1 entry. In summary, gp41 peptides on transduced cells did not alter HIV-1 receptor and co-receptor levels of expression. Meanwhile, the level of expression of these peptides on plasma membrane is an important parameter that may influence their potency to inhibit HIV-1 infection.

**Figure 2.**
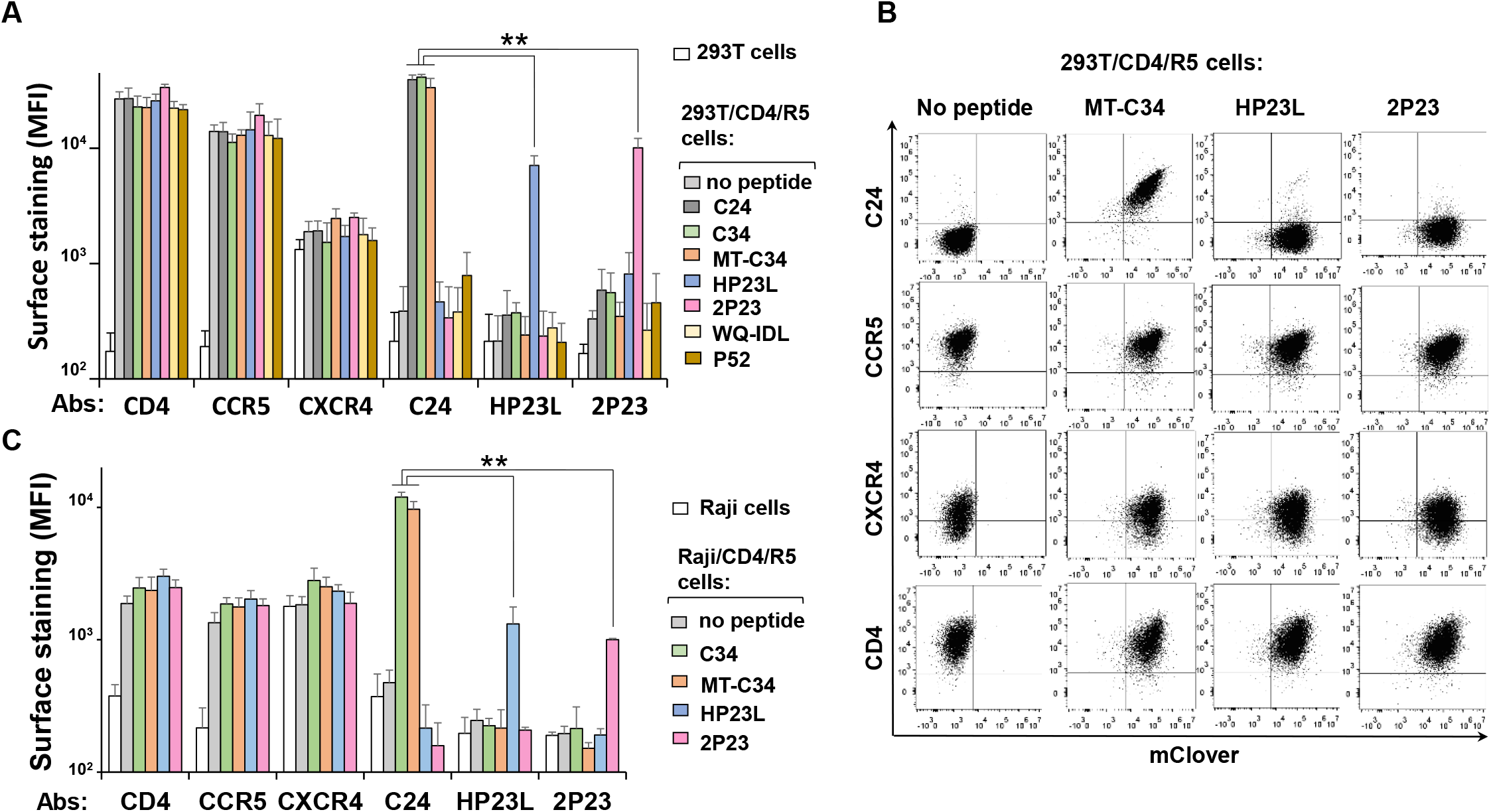
Analysis of gp41 peptide and HIV-1 receptor/coreceptor expression on surfaces of the stably transduced target cells. The 293T (**A, B)** and Raji **(C)** stably transduced with indicated peptides and selected by FACS-sorting were labeled by the indicated primary Abs, washed, and stained with the respective Alexa546-conjugated secondary Ab. Cells were analyzed using flow cytometry, and the data were presented as the average values of the mean fluorescence intensities (MFI) obtained from at least three independent measurements with standard deviations (A and C). Representative FACS graphs recorded for 293T are shown in B. ^**^ The differences are significant at p<0.01

### Modifying the CD52-based MT-C34 peptide for secretion, shedding, or binding to HIV co-receptors

We reasoned that the anti-viral activity of gp41 peptides can be expanded from gene-modified cells to bystander cells by substituting the GPI-anchored form of a peptide with a soluble one. Matabaro et al. (43) demonstrated that by regulating the fatty acid remodeling of GPI-anchored proteins, it is possible to achieve partial membrane expression of a protein needed for FACS-sorting as well as protein secretion. Another interesting approach for enhancing the activity of an anti-HIV inhibitory peptide is non-covalently linking it to a CXCR4 or CCR5 co-receptor, as demonstrated for the 15D peptide (44). We modified these approaches to adapt them for the CD52 molecular system. Specifically, to enable MT-C34 peptide secretion, we removed the GPI-attachment signal from the C-terminus of CD52 but left the leader peptide (LD) from CD52 (CD52-MT-C34s) or replaced it with LD from Gaussia luciferase (Glu-MT-C34s). Peptide shedding or binding to HIV-1 co-receptors was engineered by introducing the furin cleavage sequence RIRR (45) (MT-C34-R) or 15D (MT-C34-15D), respectively, between MT-C34 and the CD52 GPI-signal (Fig. 3A). The generated constructs were stably expressed in HIV-permissive 293T cells and tested in cell-free infection as described above. Surprisingly, both variants of secreted MT-C34 peptide failed to efficiently protect cells from HIV-1 infection, and the R and 15D modifications did not enhance the inhibitory activity of MT-C34 (Fig. 3B). We developed a sandwich ELISA system with the newly generated rabbit polyclonal and mouse monoclonal anti-C24 Abs that could detect concentrations of synthetic MT-C34 peptide as low as 5– 10 nM. At this level of sensitivity, we were unable to register secreted forms of MT-C34 peptides in the supernatants of transduced 293T cells or detect their antiviral activity in the cell-free infection test (Fig. S1A–D). This suggests that short peptides may have trouble entering the secretory pathway and require elongation or concatemerization for efficient secretion (42). The R modification is characterized by a reduced level of MT-C34 cell surface expression and decreased protection against HIV-1 infection (Fig. 3C, D), while furin inhibitor I partially elevated MT-C34-R expression level (Fig. 3D, right bars). MT-C34 expression was insensitive to Brefeldin A (BrfA) treatment (Fig. 3E, red and orange lines) but MT-C34-R, in the presence of BrfA, largely accumulated inside cells and disappeared from the cell surface (Fig. 3E, dark and light green lines). This suggests that the R modification makes peptide synthesis more dependent on the secretory pathway. Thus, cell analysis data indicate that MT-C34-R is partially cleaved off from the plasma membrane and the remaining amount of the peptide is sufficient for live cell selection and protection against HIV-1, albeit at lower levels of potency. However, the desired level of the cleaved peptide was too low to be assayed by our ELISA system (Fig. S1B) or to inhibit HIV-1 infection (Fig. S1D). In summary, the anti-HIV inhibitory activity of gp41 peptides is highest when they are GPI-anchored to the membrane. Modifications aimed at releasing the peptide from cells or additionally recruiting it to an HIV co-receptor site did not help, or even reduced, the antiviral potency of the peptide.

**Figure 3.**
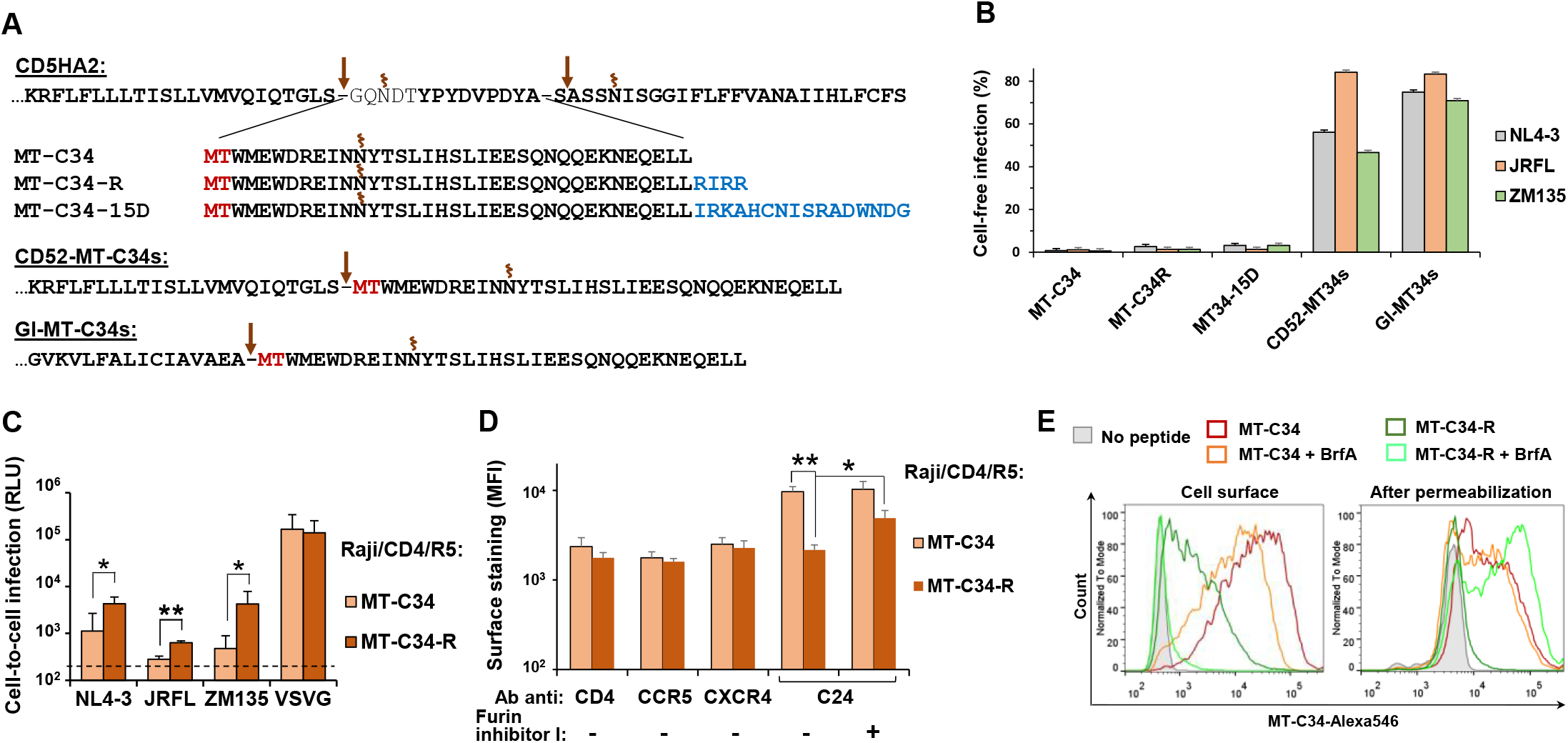
Modifying MT-C34 GPI-anchored peptide for secretion, shedding or coreceptor binding. **A**. Amino acid sequences of the generated constructs. The furin cleavage site (R) and the peptide from HIV-1 gp120 (15D) are shown in blue; see Fig.1A for the other designations. The effects of MT-C34 modifications on the levels of HIV-1 cell-free (**B**) and cell coculture (**C**) infection. The infectivity assays were set up as described for Fig.1B-C. **D**. The influence of modification R on the levels of HIV-1 receptor/coreceptors and MT-C34 expression on the surface of Raji/CD4/R5 cells. Immunofluorescent staining and analysis were performed as outlined for Fig. 2B. **E**. Comparison of the surface and intracellular expression of MT-C34 and MT-C34-R in Raji/CD4/R5 cells prior or after 16 h-treatment with brefeldin A (BrfA). Live cells were stained with anti-C24 mAb and anti-mouse Alexa546-conjugated secondary Ab (on the left), or cells were fixed and permeabilized with saponin first and then immunolabeled (on the right). Representative histogram overlays are shown. The average values with standard deviations from at least three independent experiments are shown in B-D. The differences calculated by Student *t-*test are significant at p<0.05 (*) or p<0.01 (**).

### Selecting a strategy for HIV-1 fusion inhibitory peptide knock-in

SORTS offers a quick way to test KI efficiencies at multiple sites, as it does not require the generation of a donor plasmid. We have previously applied this method to target the capsid region in HIV-1 proviral DNA (32). Here, we focused on the *CD4* and *CXCR4* HIV entry cellular genes, which have a high T cell expression, and developed strategies to knock in peptides without respective gene knockout. Specifically, we designed guide RNAs (gRNAs) and PCR donors to target the start and stop codons of the *CD4* gene and the start codon of the *CXCR4* gene. To enable transgene expression, an *in-frame* strategy with an upstream or downstream P2A sequence (32) was applied during the construction of homology arms (see Fig. 4A and Table S2 for details). To assess KI rates, we first used CD5HA2 PCR donor DNAs. CEM/R5 cells were co-transfected with Cas9 and gRNA expression plasmids and donor DNA, as previously described (32). To monitor both KI and targeted gene expression rates, cells double stained for HA and for the respective cellular protein were analyzed by flow cytometry at the indicated time post transfection. Despite a rather efficient NHEJ-mediated inactivation of *CD4* and *CXCR4* gene ORFs at the start codons, a clear population of cells with HA KI (∼6%) was detected only for the *CXCR4* gene (Fig. 4B). Importantly, HA-positive cells were also CXCR4-positive, indicating that at least one allele was precisely modified.

**Figure 4.**
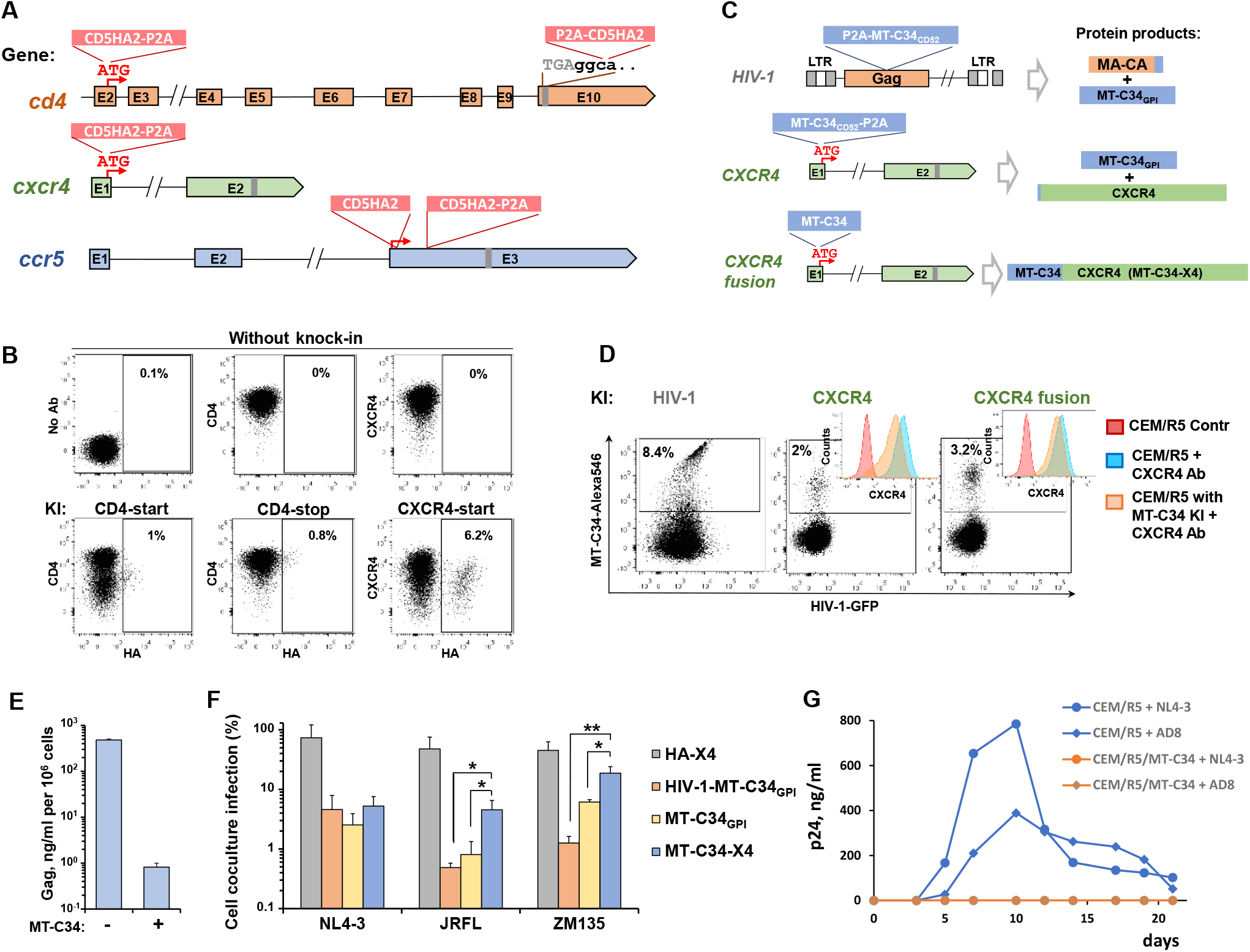
Testing different anti-HIV-1 strategies based on MT-C34 knock-in. **A**. A schematics showing cellular genes, targeted sites in exons (E), and donor DNA constructs (red boxes) selected for HA KI. **B**. The efficiencies of HA KI into the human *CD4* and *CXCR4* loci. CEM/R5 cells were electroporated to initiate KI. Three days later, transfected and control cells were double-stained for surface HA and respective cellular protein and analyzed by flow cytometry. **C**. A schematic picture illustrating genetic strategies for MT-C34 KI into the HIV-1 provirus or *CXCR4* gene (on the left) and resulting protein products (on the right). CD52 and GPI in subscript indicate that MT-C34 was embedded into CD52 sequence and expressed as GPI-anchored protein. **D**. The levels of MT-C34 and CXCR4 expression after KI in CEM/R5-HIV-1-GFPt (left Plot) and CEM/R5 (middle and right-hand graphs). Cells were immunostained for MT-C34 at 3d post transfection and analyzed by flow cytometry. MT-C34^+^ cells were sorted and labeled with anti-CXCR4 mAb along with controls (embedded histograms). Typical FACS images are shown in B and D. **E**. The levels of viral Gag produced by CEM/R5-HIV-1-GFPt cells prior (left) and after (right) MT-C34 KI and FASC sorting. **F**. Susceptibility of CEM/R5 cells with indicated KIs to HIV-1 cell coculture infection. The infectivity test was performed as described for Fig.1C, except for Raji-derived target cells were replaced with CEM cells. **G**. HIV-1 spreading assay on permissive CEM/R5 and resistant CEM/R5-MT-C34_GPI_ cells challenged with NL4-3 or NL4(AD8) viruses. The levels of HIV-1 production in E and G were quantified by p24 ELISA. The data are presented as average values with standard deviations of at least three independent experiments. * and ** - the differences are significant at p<0.05 and p<0.01, respectively, by Student *t-*test.

Following this, we replaced HA with the MT-C34 sequence in pUCHR-CD5HA2-P2A-mClover and pUCHR-mClover-P2A-CD5HA2 plasmids and PCR-amplified donor DNAs from these vectors. This allowed us to express MT-C34 from an HIV-1 provirus or a *CXCR4* gene as a GPI-anchored peptide (MT-C34_GPI_) or to fuse MT-C34 directly to the N-terminus of CXCR4 (MT-C34-X4) (31) (Fig. 4C). The FACS analysis of CEM/R5 cells, transfected for KI of these donors, showed that all strategies were effective and resulted in the parallel expression of the MT-C34 peptide and CXCR4 (Fig. 4D). The latter one, however, was expressed slightly lower on KI cells than on parental cells that can be explained by a precise modification of one allele and co-selection of the second allele with KO. The MT-C34-positive CEM/R5 and CEM/R5-HIV-1 cells from the above experiments were sorted and then tested in a cell-to-cell infection assay with three different Envs (as described above), while concomitant inactivation of HIV-1 proviral DNA in CEM/R5-HIV-1 cells was estimated by p24 ELISA. Similar to HA (see Fig. 5C in (32)), integration of MT-C34 into the HIV-1 provirus greatly reduced viral particle production (Fig. 4E). CEM/R5 cells expressing HA-CXCR4 were relatively sensitive to X4- and R5- tropic HIV-1 infection (Fig. 4F), suggesting that KI-based N-terminal modification preserved the function of CXCR4 at least at the level of viral co-receptors. Remarkably, all variants of MT-C34 peptide knocked into CXCR4 exhibited potent and comparable levels of protection against X4-tropic HIV-1 (Fig. 4F, left bars). In contrast, infection with the R5-tropic Envs JRFL and ZM-135 was more effectively inhibited by the variant MT-C34_GPI_, expressed either from proviral DNA or the *CXCR4* locus, than by MT-C34-X4 (Fig. 4F, middle and right bars). This indicates that peptide localization in lipid rafts more effectively inhibits both CXCR4 and CCR5-dependent HIV-1 fusion than its localization at the CXCR4 co-receptor site. Additionally, we performed an HIV-1 spreading assay with the molecular clones NL4-3 and NL4(AD8) and found that CEM/R5 cells bearing MT-C34_GPI_ were completely resistant to these strains of HIV-1 (Fig. 4G). In summary, we adapted SORTS for KI of the gp41 peptide MT-C34 into the *CXCR4* gene, where it can be N-terminally fused to CXCR4 or translated and expressed separately. Both modifications led to the effective and broad inhibition of HIV entry without CXCR4 knockout.

**Figure 5.**
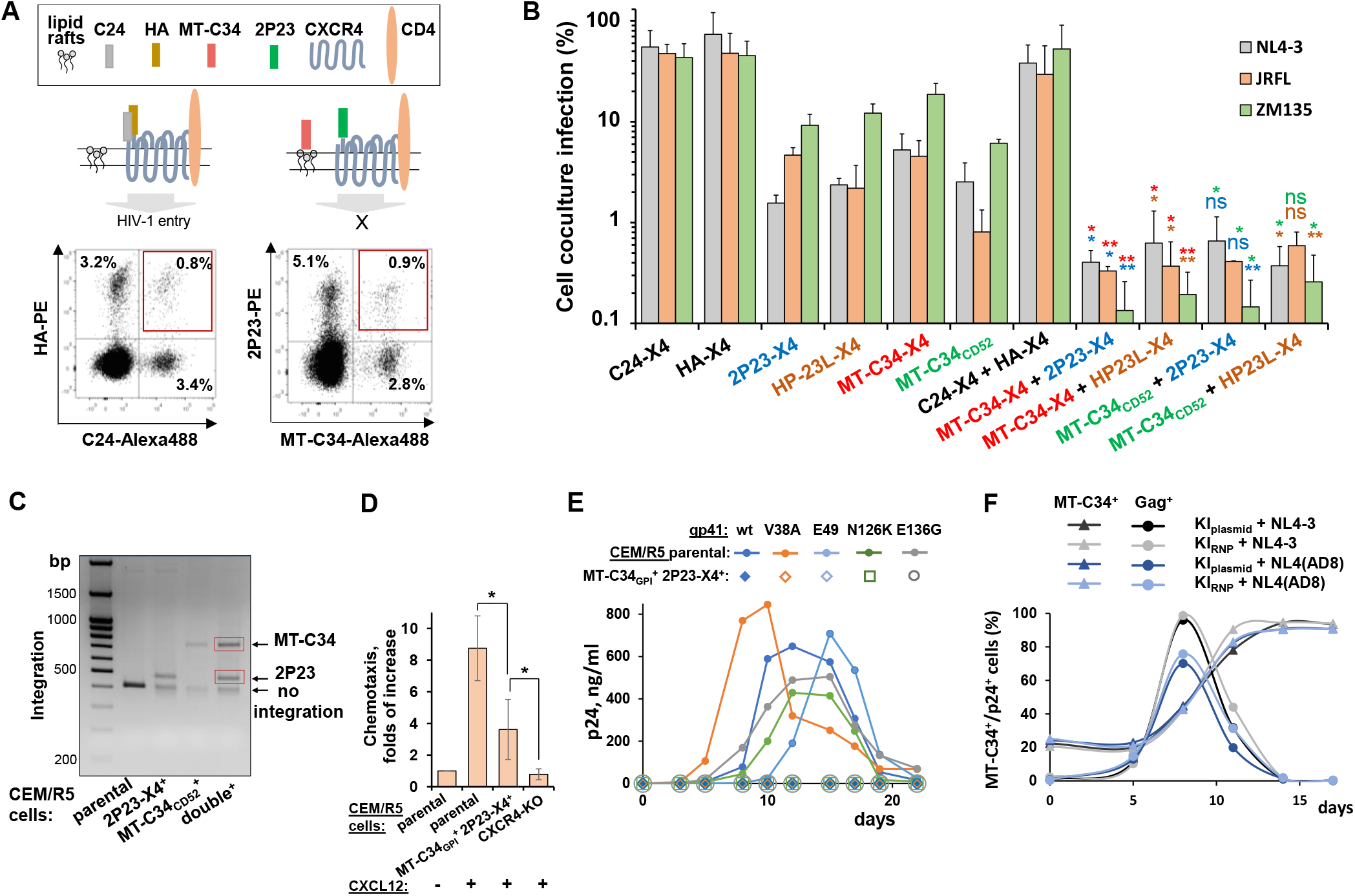
Exploring biallelic knock-in of two different peptides to straighten cell protection against HIV-1. **A**. A schematic illustrating localization of indicated peptides or epitope tags on plasma membrane after biallelic KI into *CXCR4* gene (top) and representative FACS DotPlots (bottom) recorded at day 3 after KI initiation in CEM/R5 cells. **B**. A combination of two fusion inhibitory peptide KIs enhances the level of cell protection from HIV-1 infection. CEM/R5 cells with mono- or biallelic KI of indicated peptides were isolated using respective Ab staining and FACS sorting and added as targets to transfected Raji cells to measure cell coculture infectivity levels relative to unmodified CEM/R5 control. *, ** indicate differences between double and single peptide KIs at p<0.05 and p<0.01, respectively, by Student *t-*test; to simplify understanding the groups of samples, between which the comparisons were made, the names of compared samples and resulting *p* values (asterisk) were highlighted using the same color. **C**. *CXCR4* target locus was PCR-amplified from parental or indicated sorted CEM/R5 cells, and the resulting DNA fragments were resolved on agarose gel; DNA bands indicated in red boxes were excised and analyzed by Sanger sequencing. **D**. The migration activity of CEM/R5 cells with or without indicated modifications towards the 100 ng/ml of CXCL12 (SDF-1α) chemokine. The number of spontaneously migrated cells in the control (first bar) was set at 1.0, and the number of migrated cells in the presence of chemokine was recalculated relative to the control. CXCR4-KO was generated by cotransfecting CEM/R5 cells with plasmids encoding Cas9 and gRNA targeting start codon of *CXCR4* gene followed by sorting CXCR4-negative cells. **E**. CEM/R5 cells with double KI of MT-C34 and 2P23 peptides are fully resistant to gp41 mutant HIV-1. Parental or modified cells (10^6^) were challenged with 25 ng of wt or gp41 mutant NL4-3. The kinetics of HIV-1 replication was monitored by p24 ELISA. **F**. Survival and growth advantage of MT-C34_GPI_^+^/2P23-X4^+^ cells during HIV-1 infection. CEM/R5 cells with biallelic KI generated using CRISPR/Cas9 plasmid (KI_plasmid_) or ribonucleoprotein (KI_RNP_) were mixed to permissive CEM/R5 cells at a 1:5 ratio and challenged with a high dose of NL4-3 or NL4(AD8) HIV-1 (100 ng per 10^6^ cells). Cells were double-stained for surface MT-C34 peptide and intracellular Gag to simultaneously detect the percentages of infected and HIV-1-resistant cells using flow cytometry. The data in B and D shows the average values ± std dev of three independent experiments. Representative mages and graphs are shown in A and C-E.

### Biallelic knock-in of two distinct gp41 peptides into the *CXCR4* gene synergistically enhances protection of T cells against HIV-1

As previously demonstrated, the SORTS procedure is capable of isolating a highly pure population of cells, the majority of which carry a precise gene modification in both alleles, when using two different epitope tags (32). We hypothesized that the biallelic KI of two distinct fusion inhibitory peptides would increase the cell’s level of protection against HIV-1. To explore this, we performed all possible combinations of single or dual peptide KI into the *CXCR4* gene, excluding peptide constructs that were poorly expressed (i.e., 2P23 and HP23L embedded in CD52; Fig. 2) or could not be stained simultaneously by antibodies. Following KI, CEM/R5 cells were immunostained, analyzed by FACS (see two typical examples and schematics in Fig. 5A), sorted, and tested in the cell-to-cell infection assay (Fig. 5B). As expected, the two control CEM/R5 sublines, expressing HA alone or in combination with C24 fused to CXCR4, were not substantially protected from HIV-1. The levels of inhibition mediated by MT-C34, HP23L, and 2P23 peptides fused to CXCR4 were relatively comparable and protected cells both from X4- and R5-tropic HIV-1. However, all four paired combinations of peptides that we tested exerted higher levels of HIV-1 inhibition compared to the individual peptides. From these results, we concluded that biallelic KI conveyed a protective advantage to cells.

For further analysis, we selected MT-C34_GPI_ ^+^/2P23-X4^+^ cells with two relatively distant HIV-1 fusion inhibitory peptides expressed at distinct locations on the plasma membrane. Genetic analysis of these cells showed that PCR-amplified target loci contained two DNA integration products of expected size (Fig. 5C); these were then purified from the gel and verified by sequencing (Fig. S2). Using Triton X-100 protein extraction followed by the immediate fixation and flow cytometry analysis of cell residues (46), we confirmed that the peptide MT-C34 was localized in a detergent-resistant membrane microdomain, whereas 2P23 fused to CXCR4 as well as CD4, characterized by intermediate levels of TX-100 solubilization (Fig. S3). Consistent with the findings reported by Leslie et al. (31), these engineered cells displayed a migration in the presence of CXCL12 (SDF-1α) chemokine, albeit at a lower degree compared to parental CEM/R5 cells (Fig. 5D), indicating that N-terminal modifications generated in CXCR4 molecule during KI can still support the physiological response to cognate chemokine. Next, we tested whether our combination of cell surface peptides was active against gp41 mutant HIV-1. To this end, four single mutations—V38A and E49K in the HR1 domain and N126K and E136G in the HR2 domain of gp41—that have been shown to confer HIV-1 resistance to the soluble peptides T-20, C34, and MT-C34 (47, 48) were introduced in an NL4-3 genome (Table S1). All generated mutants replicated well in CEM/R5 cells, whereas MT-C34_GPI_ ^+^/2P23-X4^+^ cells were completely protected from infection with any of the tested HIV-1 mutants (Fig. 5E). To test whether our gene-edited HIV-resistant cells would be able to grow during acute infection with HIV-1, we first optimized transfection with a Cas9 ribonucleoprotein (RNP) complex (Fig. S4A) and generated MT-C34_GPI_ ^+^/2P23-X4^+^ CEM/R5 cells using either a CRISPR/Cas9 plasmid (KI_plasmid_) or Cas9 RNP electroporation (KI_RNP_). We then set up a culture mixture that included ∼20% MT-C34_GPI_ ^+^/2P23-X4^+^ cells and ∼80% parental CEM/R5 cells and challenged them with high-dose X4- or R5-tropic HIV-1. Parental cells died following an infection peak at day eight, whereas edited cells—generated using the CRISPR/Cas9 plasmid or RNP—survived and quickly expanded over the next six days (Fig. 5F and S4B). In summary, using a CEM/R5 cell line model, we showed that it was possible to precisely modify endogenous CXCR4 and select T cells with two fusion inhibitory peptides to render cells fully resistant to wild-type or gp41 mutant HIV-1 and responsive to SDF-1α-mediated chemotaxis.

### Optimization of MT-C34 peptide knock-in into human primary cells

Precise genetic modification of primary cells is challenging. To accomplish this, we first optimized stimulation and transfection protocols for peripheral blood mononuclear cells (PBMC) by testing different methods of activation with PHA or anti-CD2, CD3, and CD28 Abs and Neon transfection protocols. The best results (60–70% GFP-positive cells) were obtained for cells activated with a combination of plastic-adsorbed anti-CD3 mAb and soluble anti-CD28 mAb or with CD2/CD3/CD28 beads for 2–3 days and electroporated using three pulses of 10 ms at 1,600 V. Thus, following optimization, transfection efficiencies of PBMC and CEM cells became comparable (Fig. 6A and B). A well-known way to improve HDR rate is to extend homology arms. By exploring KI of MT-C34 and HA into the *CXCR4* and *CD4* genes and the HIV-1 provirus in CEM and Raji cells, we demonstrated that extension of homology arms in a PCR donor from ∼100 bp to ∼500 bp increased the level of KI from 2- to 4-fold depending on the target gene. Meanwhile, the substitution of the PCR donor with a circular plasmid donor encoding 500 bp homology arms further enhanced HDR, which reached 35% at the *CXCR4* exon-2 locus (Fig. S5). After the donors were improved, we started to see MT-C34 KI into exon-2 of *CXCR4* in CD4 lymphocytes, albeit at a rate nearly 50-fold lower (∼0.6%) than in CEM cells (Fig. 6C, upper row). Using cell staining with anti-CXCR4 mAbs (Fig. 6C, low row) and a T7 endonuclease assay (Fig. 6D), we detected only a mild decrease in *CXCR4* knockout (KO) or indel formation in CD4 lymphocytes relative to those in CEM cells. Likewise, MT-C34 KI into HIV-1 proviruses with a plasmid donor was substantially lower in CD4 lymphocytes than in CEM cells (Fig. 6E and S5). Regardless of efficiency, KI of MT-C34 into proviral DNA greatly reduced the production of HIV-1 particles by CD4 lymphocytes (Fig. 6F). The obtained results suggest that the NHEJ mechanism of DNA repair may prevail during genetic modifications in human primary cells, although non-optimal experimental conditions reducing HDR cannot be excluded as well.

**Figure 6.**
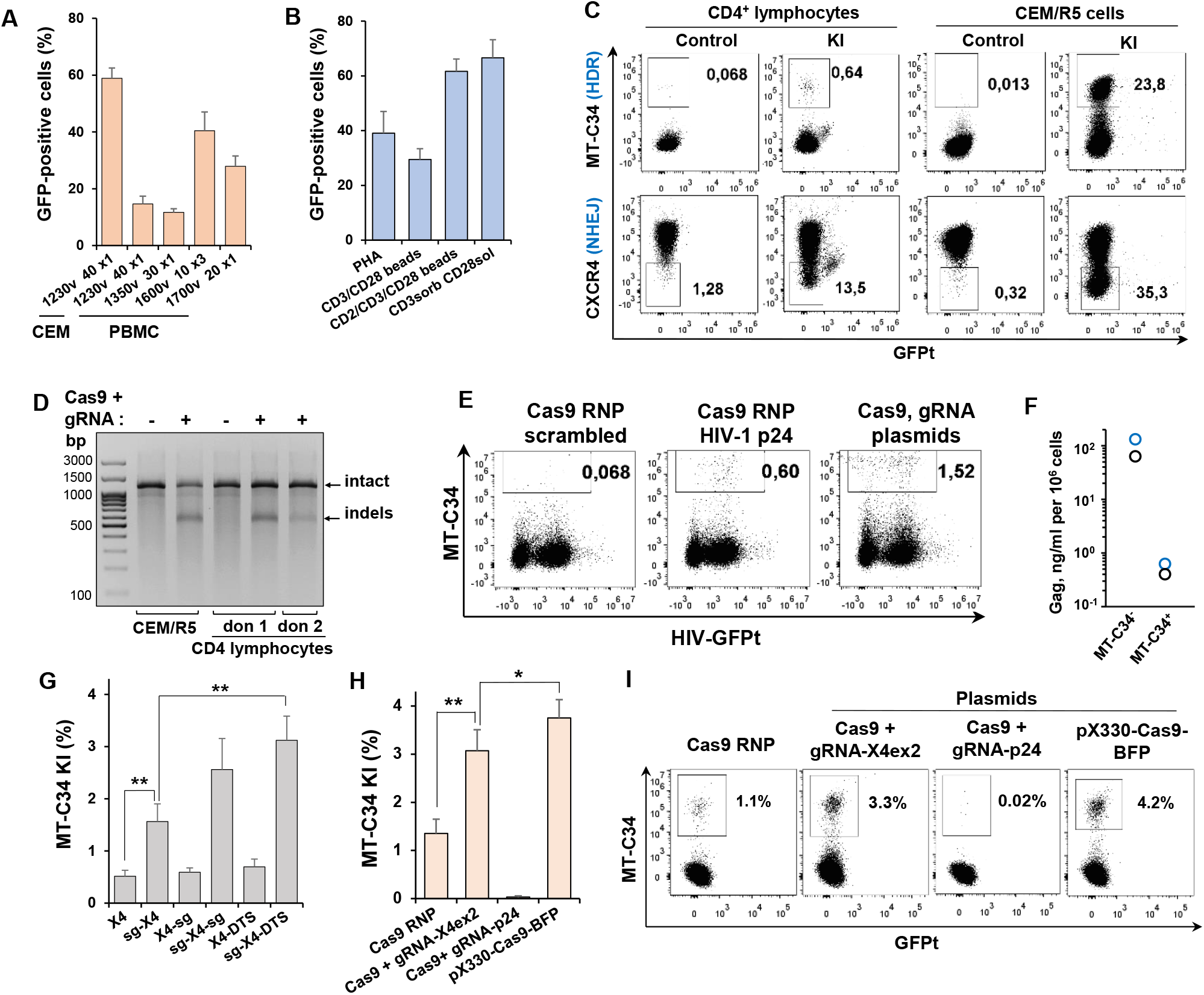
Evaluating efficiency of MT-C34 peptide KI into human primary cells. **A**. Optimization of electroporation conditions. PBMC were stimulated with 5 µg/ml PHA for 48 h and electroporated with 5 µg of GFP-expressing plasmid DNA. Transfection efficiency was measured 24 h later. **B**. Optimization of PBMC stimulation. PBMC were activated with 5 µg/ml PHA, or magnetic beads preloaded with indicated mAbs, or anti-CD3 (OKT3) mAb sorb on plastic surface with soluble 2 µg/ml anti-CD28 mAb for 48 h. Then GFP transfection efficiencies were estimated as in A. **C**. Comparison of the levels of MT-C34 KI (upper row) and CXCR4 KO (low row) in primary CD4 lymphocytes and CEM/R5 cell line. Indicated cells were co-transfected with pJet-X4 donor vector, Cas9 and gRNA expression plasmids to initiate KI into exon-2 of *CXCR4* locus. Four days later, cells were stained for MT-C34 or CXCR4 and analyzed by flow cytometry. **D**. Genetic analysis of CXCR4 exon-2 locus editing efficiency. The indicated cells were cotranscfected as described in C, but without donor. At day 3, the genomic DNAs were purified, the target locus was PCR-amplified, and the formation of indels was determined using T7 endonuclease assay. **E-F**. MT-C34 KI into HIV-1 provirus. CD4 lymphocytes activated for 2 d were infected with HIV-1-GFPt. Next day, cells were cotransfected with indicated CRISPR/Cas9 and pJet-HIV-1 p24 donor plasmid, stained for MT-C34, analyzed by flow cytometry (**E**) at day 4 post transfection and sorted into MT-C34-positive and negative populations. The equal number of sorted cells were cultured for 3 d and the Gag levels in supernatants were estimated by HIV-1 p24 ELISA. The individual values obtained for two donors are presented in F. **G**. Donor modification with protospacer and DTS sequences enhances HDR efficiency. CD4 lymphocytes were cotransfected with Cas9 and X4ex2 gRNA expression plasmids along with one of the indicated pJet donor plasmids. **H**. Type of CRISPR/Cas9 influence the level of MT34 KI. CD4 lymphocytes were electroporated with pJet-sg-X4-DTS donor plasmid and CRISPR/Cas9 in a form of RNP, two individual plasmids or a single gRNA-Cas9 expression plasmid with two NLS. Cells in G and H were analyzed at day 3 post transfection. The average values with standards deviations are shown in A-B, and G-H. I. Typical flow cytometry DotPlots corresponding to summary data presented in H. The representative FACS graphs are shown in C, E, I. ^*^ and ^**^ - the differences are significant at p<0.05 and p<0.01, respectively, by Student t-test.

It has long been known that naked transfected plasmid DNA in the cytoplasm binds to various transcription factors that transport it to the nucleus (49). We hypothesized that without a promoter our donor plasmids are poorly localized in the cell nucleus, thereby decreasing HDR. To enhance this mechanism, we added one or two X4ex2 gRNA-recognition sites (sg) (50, 51), as well as a 72-bp DNA transport signal (DTS) from the SV40 promoter (52), at the flanks of the donor sequence in the pJet-X4ex2 plasmid. These sites should have to recruit Cas9 or various transcription factors, respectively, to the DNA. Introducing sg or DTS at the 3’-position did not have a substantial effect on HDR in CD4 lymphocytes (Fig. 6G). In contrast, 5’-localization of sg, especially in combination with 3’-sg or 3’-DTS, increased the occurrence of MT-C34 KI up to 6-fold relative to KI with an unmodified plasmid donor. We then selected the best construct, pJet-sg-X4-DTS, and compared the abilities of different types of CRISPR/Cas9 to initiate KI at the *CXCR4* exon-2 locus (Fig. 6H). Cas9 RNP was slightly less efficient than transfection with gRNA and Cas9 expression plasmids. The highest level of MT-C34 KI was achieved with a single gRNA-Cas9 pX330 plasmid that contained two nuclear localization signals (NLS) instead of one NLS in the others two (Fig. 6H, right bar, and Fig. 6I).

In summary, by modifying donor DNAs and CRISPR/Cas9 we improved the KI of MT-C34_CD52_-bglpA construct into the *CXCR4* locus in human primary CD4 lymphocytes from an undetectable level up to 3.5–4.5%.

### Comparison of CD4 lymphocytes engineered for MT-C34 expression via knock-in or lentiviral transduction

We next compared the efficiencies of MT-C34 peptide delivery, the levels of MT-C34 cell surface expression and the degree of CD4 lymphocyte protection from HIV-1 infection using the CRISPR/Cas9-based KI and a standard lentiviral vector (LV) transduction. To this end, CD4 lymphocytes obtained from two donors were first activated with CD2/CD3/CD28 beads for 48 h, and then electroporated to initiate MT-C34 KI using the most optimized pJet-sg-X4-DTS donor construct and Cas9 RNP (KI_RNP_) or transduced with the LV construct pUCHR-mClover-MT-C34. Five days later, the majority of lymphocytes was immunostained and sorted into MT-C34 positive (+) and negative (-) populations. These cells were then used for measuring levels of HIV-1 single-cycle coculture infection. The unseparated cells were double-stained for MT-C34 and CD25 expression analysis. A noticeable drawback of CRISPR/Cas9/donor delivery using electroporation that we did not observe after LV transduction was the higher rates of cellular death (∼20-40%). This can be easily monitored using forward versus side scatter gating, where dead cells moved up and left relative to a live cell population (Fig. 7A). Comparing to KI, the efficiency of LV-based MT-C34 peptide delivery into CD4 lymphocytes was substantially higher (Fig.7B) and could be easily adjusted to 10-20% and more by LV titration. However, the lymphocytes expressing MT-C34 after KI displayed statistically higher level of peptide expression and can be clearly separated from peptide-negative cells in comparison to transduced cells (Fig. 7B, right panel). Since KI and transduction procedures may influence lymphocyte proliferative activity, we co-stained cells for CD25, the receptor for IL-2, which is required for ex vivo T lymphocyte proliferation. We found no changes in CD25 expression between MT-C34 positive and negative lymphocytes generated using KI or LV (Fig. 7C). An obvious advantage of KI over LV was a higher degree of CD4 lymphocyte protection from HIV-1 infection measured in cocultures with Raji effector cells. Depending on Env tropism, MT-C34 (+) CD4 lymphocytes were from 33-to 52-fold less susceptible to HIV-1 infection than MT-C34 (-) cells in the KI group of samples, but only 7-fold less sensitive upon LV transduction (Fig. 7D). This was consistent with the protective peptide expression levels detected on the surfaces of these cells (Fig. 7B).

**Figure 7.**
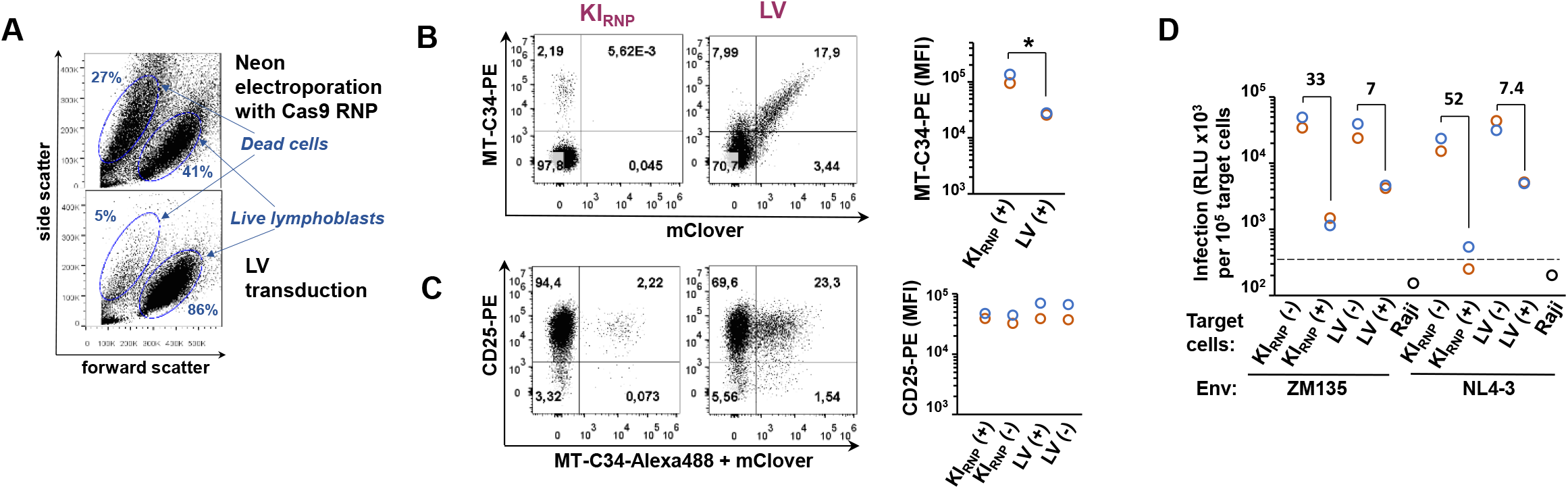
Comparative analysis of MT-C34-expressing CD4 lymphocytes generated via knock-in or lentiviral transduction. **A**. A typical live and dead CD4 lymphocyte distribution on the light scatter plots measured at day 3 post electroporation or lentiviral (LV) transduction. The levels of MT-C34 (**B**) and CD25 (**C**) cell surface expression measured at day 5 post-transfection or transduction. The flow cytometry DotPlots recorded for donor 2 and the MFI values calculated for both donors are shown on the left and right, respectively. **D**. HIV-1 single-cycle coculture infection was set up by mixing transfected HIV-1 effector Raji cells with MT-C34-positive or negative CD4 lymphocytes or Raji cells as control at a 3:1 ratio. The levels of inLuc transduction measured 3 days later were adjusted to the number of target cells and presented separately for each donor. The average values from two donors were calculated, and the folds of reduction in HIV-1 infectivity for MT-C34 (-) and (+) samples were indicated. The dashed line indicates the background level of luciferase activity. The values obtained for donors 1 and 2 are shown by orange and blue circles. KI_RNP_ stands for MT-C34 knock-in generated using Cas9 RNP. LV, lentiviral transduction as a method of MT-C34 peptide delivery. (+) and (-) indicate isolation of MT-C34 positive and negative cells. _*_ - the differences are significant at p<0.05 by Student *t*-test. The representative flow cytometry images are shown in A-C.

In conclusion, we compared two approaches for MT-C34 peptide delivery into human CD4 lymphocytes. LV offered efficient transgene expression with low cell mortality rates, whereas CRISPR/Cas9 KI into CXCR4 gene provided higher expression of MT-C34 and better protection against HIV-1.

## DISCUSSION

In this study, we adopted SORTS to engineer T cells with resistance to HIV-1 infection. First, we tested several short fusion inhibitory peptides from the HR2-domain of gp41, expressed on the cell surface using the shortest GPI-protein, CD52, and showed that some peptides (C34, MT-C34, and 2P23) can potently inhibit entry of HIV-1 independent of co-receptor usage. It is important to emphasize that these GPI-linked peptides were effective at protecting cells against not only cell-free viruses but also cell-to-cell infection, an important mechanism of HIV transmission *in vivo* (53). To demonstrate this, we employed a previously developed intron-regulated reporter system, which generates zero level of background activity (39) due to lack of active reporter protein carry over during the measurement of cell-free infection, which is often observed with conventional vectors.

Using antibodies developed against selected gp41 peptides, we demonstrated that the degree of HIV-1 inhibition depends on the level of peptide expression on the cell surface (Fig. 2). We also noticed that if a hybrid CD52-peptide molecule has two or three sites of N-glycan attachment, its surface expression is elevated compared to a hybrid with only one N-glycosylation site. We (32) and other groups (41, 42) have previously reported similar findings that suggest that N-glycosylation of GPI-anchored proteins may stabilize their expression. However, other factors such as amino acids at N-and C-terminal junctions of a peptide with the leader and GPI-anchoring signals of CD52 can also be important and require optimization for efficient peptide expression on the cell surface.

The idea of expanding protection against HIV-1 to bystander cells by genetically modifying T cells to secrete fusion inhibitory peptides has been explored previously (42, 54, 55). In our study, we aimed to achieve the partial release of an active peptide into the medium, as a portion of the peptide should remain on the plasma membrane for SORTS-mediated cell isolation. We achieved this by introducing a furin cleavage site between the C-terminus of MT-C34 and the CD52 GPI-anchoring signal (Fig. 3). This was accompanied by a decrease in MT-C34-R cell surface expression and protection against HIV-1 that could be partially restored by furin inhibitor I. In addition, the conditioned medium from these cells did not inhibit HIV-1 infection. Based on the MT-C34 ELISA results, we concluded that this inefficient protection resulted from the low level of MT-C34 peptide in the supernatant (Fig. S1). The problem of low anti-HIV peptide secretion by genetically modified T cells has been previously discussed (42); one way to overcome difficulties related to inefficient entry of a short peptide into the secretory pathway is to elongate or concatemerize these peptides.

Third-generation lentiviral vectors have become safer and have been used to protect T cells from HIV-1 infection by transducing different antiviral genes such as modified restriction factors APOBEC3G-D128K (56) and TRIM5α-CypA (57), CCR5-interfering RNAs (58), fusion inhibitory peptides (C34-CXCR4 clinical trial), or a combination of sh5 and C46 (59). Although they can efficiently deliver relatively large genes into primary cells without producing double-strand breaks in the host’s genomic DNA, several problems remain. These issues include the potential risk of insertional mutagenesis, the silencing of an integrated transgene over time, an immune response to selection markers (fluorescent proteins or antibiotic resistance factor), and the concomitant expression of an endogenous gene that can interfere with a therapeutic gene and reduce its effect. In our study, we attempted CRISPR/Cas9 KI of preselected gp41 peptides into the HIV-1 entry genes *CD4* and *CXCR4*. We succeeded in targeting the *CXCR4* gene at the start codon (exon-1) or the beginning of exon-2. The first strategy allowed us to save CXCR4 expression and concomitantly obtain biallelic KI of two different fusion inhibitory peptides, one in the context of the CD52 molecule (i.e., GPI-anchored) and the other fused in frame with the N-terminus of CXCR4, as previously explored by Leslie et al. (31). We showed that the combination of these two peptides provided CEM/R5 cells with stronger protection against HIV than individual peptides. These cells could survive and grow during acute HIV-1 infection and were fully protected from all tested gp41 mutants of HIV-1. Whether HIV-1 can evolve resistance to both the MT-C34 and the 2P23 peptides anchored at the plasma membrane remains unknown. He’s group reported mutations in gp41 conferring resistance of SIV to the LP-52 (60), the lipid version of P52, which did not show noticeable protection in our HIV study, but, in the discussion of another paper, the same group mentioned their inability to select HIV-1 mutants escaping the 2P23 inhibitor or its lipid derivative LP-19 (30). Accordingly, our extended monitoring of CEM/R5/MT-C34^+^/2P23^+^ cell culture challenged with HIV-1 or HIV-1-infected CEM/R5 cells did not reveal any signs of viral rebound (unpublished observations). This suggests that a combination of these peptides may represent a high genetic barrier to induction of resistance, although more work is needed to confirm it.

Knowing that primary cells are difficult targets for HDR-based gene editing, we first optimized the PBMC activation/transfection protocol to obtain 60–70% efficiency. Second, through the extension of homology arms from 100 to 500 bp and substitution of the PCR donor with a circular plasmid, we reached unusually high levels of HDR in CEM cells (∼25–35%) when targeting exon-2 of CXCR4 or HIV-1 capsid (Fig. S5A and C). This can logically be explained by the more efficient nuclear entry of plasmid supercoiled DNA than of linear PCR fragments during electroporation, though some other possible explanations may exist. Søndergaard et al. reported that small plasmid cotransfection increased levels of CRISPR/Cas9 delivery and cell survival (61). Hornstein et al. showed that increasing vector length to 1.9 kb enhanced transfection, which then proceeded to plateau as plasmids further increased in size (up to 4.5 kb) (62). Nevertheless, despite fairly efficient locus editing, primary human lymphocytes are characterized by low HDR. One of many possible explanations for this is that, unlike immortalized cell lines, DNA repair template, as well as Cas9 may have difficulties entering the nucleus of a primary cell. This assumption is supported by our data showing that SV40 DTS and gRNA target sites placed at the flanks of the donor and additional NLS in Cas9 improved HDR in CD4 lymphocytes (Fig 6 G-I), but not in the CEM cell line, where HDR level was high before the donor plasmid modification (unpublished data).

Using the side-by-side comparison of LV and HDR-based delivery of MT-C34 peptide to the human CD4 lymphocytes, we highlighted several pros and cons of the two technologies. LV transduction mediated fairly efficient peptide delivery with low cell toxicity, but, due to nearly random transgene integration, the peptide was expressed at different levels. Its average level of expression was significantly lower than that measured in knocked-in cells. This can be explained by the high activity of the *CXCR4* promoter in stimulated lymphocytes and/or by a smaller size of the peptide-coding mRNA transcribed from the edited *CXCR4* locus in comparison to LV-driven mRNA, which encodes both the peptide and mClover. We cannot exclude that CMV promoter used in our LV may also influence the transgene expression, and that its substitution with EF1a or PGK promoter can determine the role of promoter in MT-C34 expression. Consequently, the degree of lymphocyte protection from HIV-1 was higher with KI, than with LV. From our point of view, the positive aspects of each technology could be used for future improvements. For instance, to avoid electroporation-induced cell death, the usage of LV particles for Cas9 RNP delivery looks very attractive. To date, several Cas9-VLP technologies have been developed that utilize retroviral (63) or lentiviral (64, 65) Gag assembly. Although they demonstrated a good efficiency for gene knockout, VLPs specifically recruiting repair template have not been devised. Also, the reported high doses of VLPs required for gene editing indicate that the system may not be optimal for hybrid molecule assembly which may result from their different subcellular localization, cytoplasmic for Gag and nuclear for Cas9 and gRNA. Solving these problems will greatly propel studies on HDR-based therapeutic technologies.

In conclusion, we developed a gp41 fusion inhibitory peptide CRISPR/Cas9 HDR-based platform to generate T cells with high and broad resistance to HIV-1 infection. For the first time, we demonstrated the possibility of using this platform to target the *CXCR4* locus and HIV provirus in human primary CD4 lymphocytes. We believe that further improvements to HDR efficiency in primary cells are possible and, together with the system adaptation to the *CCR5* locus, the described CRISPR/Cas9 peptide KI platform will be a powerful instrument among the existing gene-based therapeutics developed to combat HIV.

## METHODS

### Cell cultures

The human embryonic kidney 293T cells and HeLa-TZM-blue reporter cells were obtained through NIH AIDS Research and Reference Reagent Program. The human CD4 T cell line CCRF-CEM and the B cell line Raji were purchased from ATCC. The peripheral blood mononuclear cells (PBMC) from fresh donor blood or buffy coat were isolated on the density gradient of Ficoll-Paque (Paneco, Russia). The CD4^+^ lymphocytes were isolated using Dynabeads Untouched Kit (Invitrogen, USA). The purified PBMC and CD4 lymphocytes were activated with 5 μg/ml of phytohemaglutinin (PHA) (Sigma-Aldrich, USA), CD2/CD3/CD28 activation beads (Miltenyi Biotec, Germany) or plastic-adhered anti-CD3 mAb plus soluble anti-CD28 mAb for 2-3 days, and grown in the presence of 100 U/ml of recombinant human interleukin-2 (Ronkoleikin, Biotech, Russia). The 293T cell line was cultured in high glucose Dulbecco’s modified Eagle’s medium (DMEM) (Sigma-Aldrich, USA) with sodium pyruvate, sodium bicarbonate, 10% fetal calf serum (FCS), 2 mM glutamine and 40 µg/ml gentamicin. PBMCs, CEM and Raji cells were maintained in RPMI 1640 medium containing 10% fetal calf serum, 2 mM glutamine, 50 µM 2-beta-mercaptoethanol, and 40 µg/ml gentamicin. All transfections were performed on low passage cells; cultured cell lines were checked periodically for mycoplasma contamination.

### Plasmid construction

To create vectors for stable expression of HIV-1 fusion inhibitory peptides, the HA epitope with or without upstream GQNDT from the CD52 coding sequence in pUCHR-mClover-smAID-P2A-CD5HA2 or pUCHR-CD5HA2-P2A-smAID-mClover lentiviral plasmid (32) was substituted with the respective peptide coding sequence by using overlapping extension PCR followed by cloning at Afe I/PspX I or Afe I/Age I restriction sites, respectively. The pUCHR-CCR5 plasmid was constructed by PCR amplification of CCR5 coding sequence from U937 cell-derived cDNA and subsequent cloning the resulting DNA at EcoR I and Xma I restriction sites into pUCHR vector. The HIV-1 molecular clones pNL4-3 and pNL(AD8) were kindly provided by Dr. Eric Freed (NCI-Frederick, USA); CH077 clade B and K3016 clade C transmitted-founder viruses were obtained through NIH AIDS Research and Reference Reagent Program. For generation of NL4-3 mutants in gp41 region of Env, the BsaBI/BamH I DNA fragment from pNL4-3 plasmid was amplified by fusion PCR using overlapping primers with a mutation. It was, first, cloned into pJet plasmid (Thermo Scientific, USA), and a mutation was verified by DNA sequencing; then the fragment was cloned back into pNL4-3 using aforementioned restriction sites. The detailed information about primers and generated constructs is presented in Table S1. All PCR DNA fragments prepared for cloning were generated using Pfu polymerase (Sibenzyme, Russia) and verified by sequencing. The gRNA expressing vector pKS gRNA BB was described earlier (66). Plasmids for the expression of wild type spCas9 alone or in combination with gRNA were pcDNA3.3-hCas9 (Addgene #41815) and pX330-T2A-BFP (Addgene # 64216), respectively.

### Lentiviral transduction

To generate lentiviral particles, 8 × 10^5^ 293 T cells in a 6 cm dish were cotransfected with 2 μg of HIV-1 packaging plasmid pCMVΔ8.2 R (Addgene # 12263); 0.5 μg of the pCMV-VSV-G plasmid (Addgene # 8454) expressing protein G from vesicular stomatitis virus (VSV-G); and 3 μg of one of the pUCHR transfer vector constructed as described in previous paragraph. Transfection was performed for 6 h using Lipofectamine 2000 reagent (Thermo Scientific, USA) according to the manufacturer’s instruction, then the culture medium was replaced. Supernatants containing PVs were harvested and cleared through 0.45 μm pore size filters at 54 h posttransfection. The 293T/CD4 or Raji/CD4 cells that we described previously (67) were, first, infected with PVs to express CCR5. After immunostaining with respective mAb and positive sorting, cells were transduced to express one of gp41 peptides. Peptide-expressing cells were then sorted based on mClover fluorescence. Transductions were set up using different doses of PVs. Two-three days postinfection the levels of transduction were quantified by flow cytometry, and samples with less than 30% of positive events (low MOI) were selected for sorting.

### Cas9, guide RNAs and donor DNAs

Cas9 protein was produced in *E*.*coli* BL21 (DE3) strain after transformation with SP-Cas9 plasmid (a gift from Niels Geijsen, Addgene plasmid # 62731) and purified by metal chelate affinity chromatography on Ni-NTA agarose (Qiagen, USA), followed by cation exchange chromatography and gel filtration, and stored at -80°C at a concentration of 6.27 µg/µl (38.93 pM/µl). Protospacer sequences for guide RNAs (gRNAs) were selected using two web-based resources https://www.benchling.com/crispr/ and http://chopchop.cbu.uib.no/ and cloned into pKS gRNA BB or pX330-T2A-BFP plasmid using BbsI restriction site. Guide RNAs were transcribed *in vitro* from PCR templates using MEGAshortscript High Yield Transcription Kit (Ambion, Thermo Fisher Scientific, USA) in accordance to manufacturer’s protocol and purified from the reaction mix with miRNeasy kit (Qiagen, Germany). To prepare PCR template for an *in vitro* transcription reaction, the gRNA sequence was PCR-amplified from the respective pKS-gRNA plasmid using 5’-primer bearing T7 promoter sequence TGTAATACGACTCACTATAGGG and a specific protospacer sequence and 3’-U6 terminator primer AAAAGCACCGACTCGGTGCC. PCR products were purified and used in an *in vitro* transcription reaction at 150 nM concentration. Donor DNAs for SORTS were prepared by PCR, as described earlier (32), i.e., with primers designed to include ∼100 nt homology arms in a close proximity to DNA cut site and with respective peptide expression plasmid, used as a template. PCR products were run on 1% agarose gel and purified using gel extraction Kit (Thermo Scientific). To generate donor plasmids, ∼400-500 bp homology arms were amplified from genomic DNA isolated from PBMC and fused with a respective short PCR-donor DNA to get a long PCR donor. The latter was cloned into pJet plasmid and verified by sequencing. The modifications of pJet donor plasmids with protospacers and DTS were performed by cloning annealed oligos at the respective restriction sites that flank a donor sequence. The detailed information about gRNAs, primers and templates used for donor generation is presented in Table S2. Two plasmids pUCHR-mClover-MT-C34 and pJet-gR-X4ex2-DTS were deposited at Addgene under the accession numbers 117153 and 117154, respectively.

### Knock-ins and SORTS

For SORTS principle and experimental protocol, please, refer to our previous paper (32). Briefly, to initiate knock-in (KI) of gp41 peptide into proviral DNA or cellular gene, 1-2×10^6^ CEM, CEM-HIV-1-GFPt cells, activated human PBMC or CD4^+^ lymphocytes were cotransfected with 3 μg of pcDNA3.3-hCas9, 1 μg of pKS-gRNA plasmid DNAs, and 2 pM of respective PCR or plasmid donor/s. Alternatively, Cas9 and gRNA expression plasmids were substituted with 4 μg of pX330-Cas9-BFP plasmid or 200 pM Cas9 RNP. For RNP preparation, Cas9 protein and gRNA were mixed at equimolar concentration in Buffer R and incubated 20-30 min before electroporation. All cell types were transfected using Neon electroporation system with 100 µl pipet tips (Invitrogen, USA) at the following settings 1,230 V, 40 ms × 1 for CEM cells, and 1,600 V, 10 ms x 3 for human primary cells. At day 3 posttransfection, cells were immunostained with respective Abs, and the level of KI as a percentage of positively labeled cells was assessed by flow cytometry. Finally, cells expressing peptide/s were isolated using several rounds of FACS sorting procedure until high level of purity was achieved.

### Genetic analysis of targeted loci

Genomic DNA from ∼10^6^ parental or sorted CEM/R5 cells or CD4 lymphocytes was purified using Quick-gDNA MiniPrep Kit (Zymo Research, USA). The *CXCR4* target regions were PCR-amplified using 200 ng of genomic DNA, Taq DNA polymerase (SibEnzyme, Russia) and primers listed at the end of Table S2. PCR reaction was set up using the following parameters: (95 °C-2′) × 1, (95 °C-30′′, 65 °C-30′′, 72 °C-30′′) × 30, (72 °C-5′) × 1 The DNA fragments amplified from double positive cells and containing integrated transgene were resolved on agarose gel, purified and cloned into pJet 1.2 PCR cloning vector (Thermo Scientific, USA). The DNAs from single bacterial clones were sequenced using Sanger method. The level of indels formation in PCR-amplified target locus was analyzed by T7 endonuclease assay using standard protocol.

### Antibody generation and purification

Synthetic peptides C24, MT-C34, HP23L, and 2P23 (see sequences in Fig.1A) with or without an additional C-terminal cysteine residue were ordered from Syneuro company (Moscow, Russia). Each cysteine-modified peptide was conjugated to two carrier proteins, bovine serum albumin (BSA) and keyhole limpet hemocyanin (KLH) (both from Sigma-Aldrich, USA) using sulfosuccinimidyl 4-(N-maleimidomethyl)cyclohexane-1-carboxylate reagent (ThermoFisher Scientific, USA) in accordance to manufacturer’s protocol. Total four 8-week-old female BALB/c mice and two New Zealand rabbits (per antigen) were immunized three times every 2 weeks with a BSA-conjugated peptide injected intraperitoneally (mice) or subcutaneously (rabbits) in complete (first shot) and incomplete (shots 2 and 3) Freund’s adjuvant. At day 35, test bleeds were performed, and sera from individual animals were tested by ELISA with KLH-conjugated peptide immobilized on plastic surface and anti-mouse or anti-rabbit horse radish peroxidase (HRP)-conjugated detector Ab (Cell Signaling Technology, USA). The animals with the highest Ab titer were selected for booster immunization. Four days after the final immunization, mice were sacrificed, and spleen lymphocytes were hybridized with the Sp2/0 myeloma cell line, and cultivated on a monolayer of feeder cells (mouse peritoneal macrophages) for about 2 weeks. The first screening of hybridoma supernatants was performed by using ELISA with KLH-conjugated peptide. The positive hybridomas were then screened on peptide-expressing 293T cells using immunofluorescence (IF) and flow cytometry. In a week after booster shot, ∼ 50 ml of sera from rabbits were collected and analyzed as outlined above for mice. The selected mouse and rabbit immunoglobulins were affinity purified on protein G SepFast resins (BioToolmics, UK) or on peptide used for immunization and conjugated to a cyanogen bromide-activated-Sepharose® 4B (Sigma-Aldrich, USA), respectively, and in accordance to manufacturer’s protocols.

### Triton X-100 solubilization assay

The sensitivity of cellular antigens and peptides to TX-100 solubilization using flow cytometry was performed as essentially described by Filatov A. et al. (46). Briefly, 1×10^7^ cells were washed in PBS, lysed in 1 ml ice-cold 1% TX-100 in PBS supplemented with 1 mM phenylmethylsulfonyl fluoride (PMSF) for 30 min at 4 °C, and fixed by diluting sample with 10 ml of ice-cold 1% paraformaldehyde (Sigma-Aldrich, USA) in PBS and incubating for additional 30 min . TX-100 extracted cell residues were collected by centrifugation at 1700 x g for 10 min, stained with primary and secondary Abs and analyzed by flow cytometry as described below. To avoid losing cellular remains, the smaller volume of reaction mixture and the higher centrifuge forces (see above) were applied than those used for live cell labeling. For better visualization of live and TX-100 extracted cells on flow cytometer, the axis scales on forward/side scatter plot were turned from linear to logarithmic.

### Immunofluorescence, flow cytometry and cell sorting

Cell surface antigen immunolabeling was performed on live cells by incubating them with a primary Ab for 20 min., followed by two washes with PBS, incubation with an appropriate secondary fluorophore-conjugated Ab for 20 min and final washes with PBS. Prior addition, all Ab were diluted in PBS supplemented with 2% FCS at a final concentration of 5 µg/ml. For intracellular staining, cells were fixed with 4% paraformaldehyde (Sigma-Aldrich, USA) in PBS for 15 min, washed with PBS, and incubated for 30 min in permeabilization solution containing 0.1% saponin (Sigma-Aldrich, USA) and 2% normal mouse serum in PBS. HIV-1 Gag was stained by addition of rhodamine-labeled anti-p24 mAb at 1:200 dilution for 30-40 min. Cells were washed 3 time with permeabilization solution and resuspended in PBS prior analysis. The list of primary commercial Ab used in this study included: the rabbit anti-HA mAb clone C29F4 (Cell Signaling Technology, USA), the mouse anti-human mAbs CD3 clone OKT3, CD4 clone EM4, CD45 clone LT45, CD71 clone LT-4E3, CD98 clone MC7E7 (all from Sorbent, Moscow, Russia), CD59 clone MEM43 (Exbio, Czech Republic), CXCR4 clone 12G5 (Santa Cruz Biotechnology, Dallas, TX, USA), anti-human CD195 clone 2D7/CCR5 (BD Pharmingen™, USA), and anti-HIV-1 p24 clone KC57-RD1 (Beckman Coulter, CA, USA). The secondary goat anti-mouse and anti-rabbit Abs conjugated to Alexa488, Alexa546 or phycoerythrin (PE) were purchased from Thermo Scientific, USA. Immunolabeled samples were analyzed on CytoFLEX S (Beckman Coulter, USA) flow cytometry instrument equipped with 488-nm and 561-nm lasers, which were used to detect Alexa488 (or GFPt) and PE (or Alexa546) fluorescent signals, respectively. Cells were sorted using a FACSAria II (Becton Dickinson Biosciences, San Jose, CA, USA) or Sony MA900 (Sony Biotechnology, San Jose, CA, USA) Instrument. The collected data were analyzed by CytExpert and presented using FlowJo LLC software.

### HIV-1 cell-free and cell coculture infections

For HIV-1 single-round cell-coculture and cell-free infection tests with intron-regulated reporter vector inLuc, please, refer to our previous papers (39, 40, 67). Briefly, to generated viral particles pseudotyped with different Envs, 293T cells in 10-cm dish were cotransfected with 4 μg of pCMVΔ8.2 R plasmid, 6 μg of pUCHR-inLuc-mR reporter vector, and 1 μg of one of the HIV-1 Env expressing vectors pIIINL4env (a gift from Dr. Eric Freed, NCI-Frederick, USA), pJRFLenv (68) and pZM135 (a gift from Drs. E. Hunter and J. Blackwell, Emory University, GA USA). At 48 h posttransfection, supernatants containing PVs were harvested, cleared through filters with 0.45 μm pores, and the levels of HIV-1 Gag in supernatants were quantified by p24 ELISA. To initiate cell-free infection, 7.5 × 10^4^ 293T/CD4/R5 cells with or without peptide in a 24-well plate were infected with an equal amount of PVs (∼2 µg of p24 per well) for 48 h. To set up cell-coculture infection, 10^6^ Raji cells were Neon electroporated (1,350 V, 30 ms × 1) with the same viral vectors as above in the amount of 2 μg, 3 μg and 1 μg, respectively. Next day, transfected Raji cells were directly mixed with Raji/CD4/R5 targets cells with or without peptide expression at a 1:1 ratio, and cultured in 5 ml of complete growth medium for 48 h. To measure cell coculture infection in primary cells, transfected Raji cells were mixed with CD4 lymphocytes at a 3:1 ratio and incubated for 3 days. At 16 h prior harvesting, 20 nM of phorbol 12-myristate 13-acetate (PMA) was added into co-cultures to enhance reporter expression in infected cells. The luciferase activity as a measure of infectivity was estimated using the GLO lysis buffer, Bright-GLO luciferase assay system, and GloMax®-Multi Jr Single-Tube Multimode Reader (all from Promega, Madison, WI, USA).

### HIV-1 spreading assay

A panel of infectious HIV-1 (NL4-3 wt and mutants, NL(AD8) was generated by transfecting HEK293T cells with a respective molecular clone as described above. Virus input quantities were normalized for inoculation by measuring p24 levels and in some cases by determining 50% tissue culture infectious dose (TCID_50_) on TZM-bl cells, as described (69). Total 10^6^ CEM/R5 cells were challenged with a high dose of virus (25-100 ng of p24 or ∼ 100-400 TCID_50_) in 200 µl of growth medium for 5 h, then washed of the virus three times and placed in 1 ml of complete growth medium. Every 2-3 days, the 1/2-2/3 of the resuspended cell culture was removed and replaced with a fresh medium. The cells in collected probed were pelleted by centrifugation and, where appropriate, double-stained for surface MT-C34 peptide and intracellular HIV-1 p24, while supernatants were stored in a freezer for subsequent p24 ELISA measurements. Dynamics of HIV-1 replication was monitored at least for 21 days.

### Chemotaxis assay

The migration of CEM/R5 cells was evaluated using a transwell chamber with 8-μm pores (SPL). Lower wells were filled with 500 μl of migration medium (RPMI, 4 mM glutamine, 10 μg/ml gentamicin, 0.3% BSA, 10 mM HEPES pH 7.2-7.4) in the presence or absence of 100 ng/ml SDF-1α (Abcam #9798, Cambridge, UK). Cells (1 × 10^6^ cells/well) were seeded on 24-well transwell inserts in 100 μl of migration medium without a chemokine. Cells were allowed to migrate through a transwell for 4 h in CO_2_-incubator at 37°C. Migrated cells were collected from the lower chamber and counted using a flow-count option of Cytoflex S. The number of migrated cells in the presence of chemokine was divided by the number of spontaneously migrated cells to calculate the degree of chemotaxis.

### ELISA

The levels of HIV-1 core protein Gag in cell supernatants were quantified using HIV-1 p24 ELISA Kit (Zeptometrix, Bufalo NY, USA) in accordance to the manufacturer’s instruction. For hybridoma screening, the 96-well ELISA plates (SPL, Life Sciences, Korea) were coated by a conjugate of immunogenic peptide with KLH diluted in PBS at 5 µg/ml concentration. All subsequent steps, i.e. washing, blocking, adding hybridoma supernatants, and detecting with the secondary HRP-conjugated Ab were performed using ELISA commercial solutions and diluents purchased from Xema-Medica Co (Russia). The levels of peptides in supernatants were assessed by a sandwich ELISA with a mouse monoclonal Ab against peptide acting as a capture reagent, and a rabbit polyclonal Ab raised against the same peptide together with a secondary anti-rabbit HRP-conjugated Ab used for detection purpose. Prior addition, all Abs were diluted to a final concentration of 1 µg/ml.

### Data availability statement

The data that support the findings of this study, some reagents generated in this study (antibodies and plasmids) are available from the corresponding author upon reasonable request.

### Ethic statement

All experiments that involved human blood samples and laboratory animals were approved by the Human and Animal Ethics Committees of the Institute of Gene Biology (Moscow). The animals used in this study were handled in strict accordance with the recommendations in the Guide for the Care and Use of Laboratory Animals of the National Institutes of Health (USA). Blood donors gave informed consent for the use of their samples in the described experiments. All methods were performed in accordance with relevant the guidelines and regulations.

## ACKNOWLEGEMENTS

We thank Alexey Deykin and Evgeniy Korshunov from Core Facility Center of the Institute of Gene Biology of Russian Academy of Science (IBG RAS) (Moscow) for laboratory animal handling and immunization; Olga Arkova and Artem Bonchuk from the Center for Precision Genome Editing and Genetic Technologies for Biomedicine of IBG RAS for Cas9 protein purification. We are grateful to Eric Freed from NCI-Frederick for the critical reading of the manuscript. This work was mostly supported by the grant 18-29-07052 from Russian Foundation for Basic Research and partially (studies on primary cells) by the grant 075-15-2019-1661 from the Ministry of Science and Higher Education of the Russian Federation.

## AUTHOR CONTRIBUTIONS

D.M. – concept development, construct design, data analysis, antibody generation, writing manuscript; A.M. – lentiviral plasmid construction, cell transduction, single-round infection tests, immunofluorescent analysis; N.K. – primer design, donor DNA preparation, primary cell activation and transfections, knock-in, FACS analysis; S.K. – hybridoma generation and screening, antibody purification, genetic analysis; D.K. – cell sorting, ELISA; M.S. – guide RNA and Cas9 RNP preparation, KI analysis, editing manuscript; D.G. – TX-100 antigen solubilization, immunofluorescence, flow cytometry analysis; A.S and A.V. – HIV spreading assay, virus titration, data analysis, editing manuscript; A.F. – monoclonal hybridoma generation, data analysis.

## COMPETING INTERESTS

The authors declare no competing financial and non-financial interests.

## MATERIALS & CORRESPONDENCE

All correspondence and material requests should be addressed to

## SUPPLEMENTARY INFORMATION

**Figure S1.**
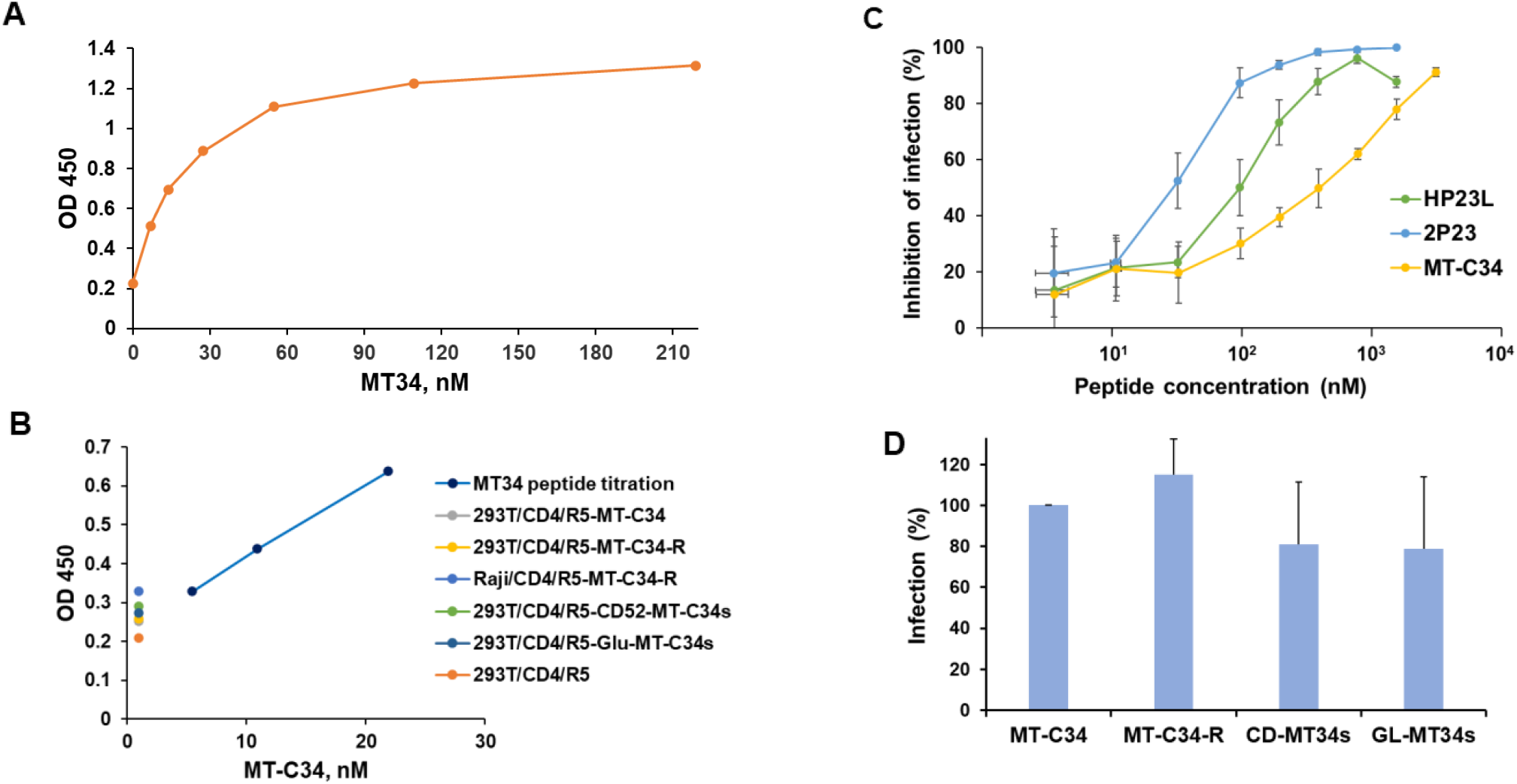
Secreted or shed forms of MT-C34 peptide are not detected by sandwich ELISA and do not inhibit HIV-1 infection. **A**. Determining sensitivity of developed MT-C34 ELISA with a pair of rabbit and mouse anti-MT-C34 Abs. The synthetic peptide MT-C34 was titrated and assayed by ELISA. **B**. The levels of MT-C34 peptide in cell supernatants are below the detection limit of the established sandwich ELISA. The indicated cells were passaged and cultured for 48 h, then the supernatants were harvested and assayed by ELISA without dilution. In parallel, the control MT-C34 peptide at several dilutions was evaluated in the same plate. **C**. Inhibitory activities of synthetic peptides against HIV-1_JRFL_. VLPs psedotyped with JRFL Env were preincubated with one of the indicated peptides for 1h and the levels of infectivity were measured on 293T/CD4/R5 cells as described in Methods. The results are presented as an inhibitory activity relative to that with no peptide. **D**. Supernatants from 293T cells engineered to secrete or shed MT-C34 peptide do not inhibit HIV-1 infection. VLPs with JRFL Env were preincubated for 1 h with 0.25 ml of the supernatant derived from one of the indicated modified 293T cells and added to 293T/CD4/R5 cells cultured in 24-well plate in 0.25 ml of growth medium. The levels of infectivity were quantified as in C and presented relative to MT-C34 sample that does not contain released peptide. The graphs in A and B are representative of two separate experiments. The average values of at least three independent experiments with standard deviations are shown in C and D

**Figure S2.**
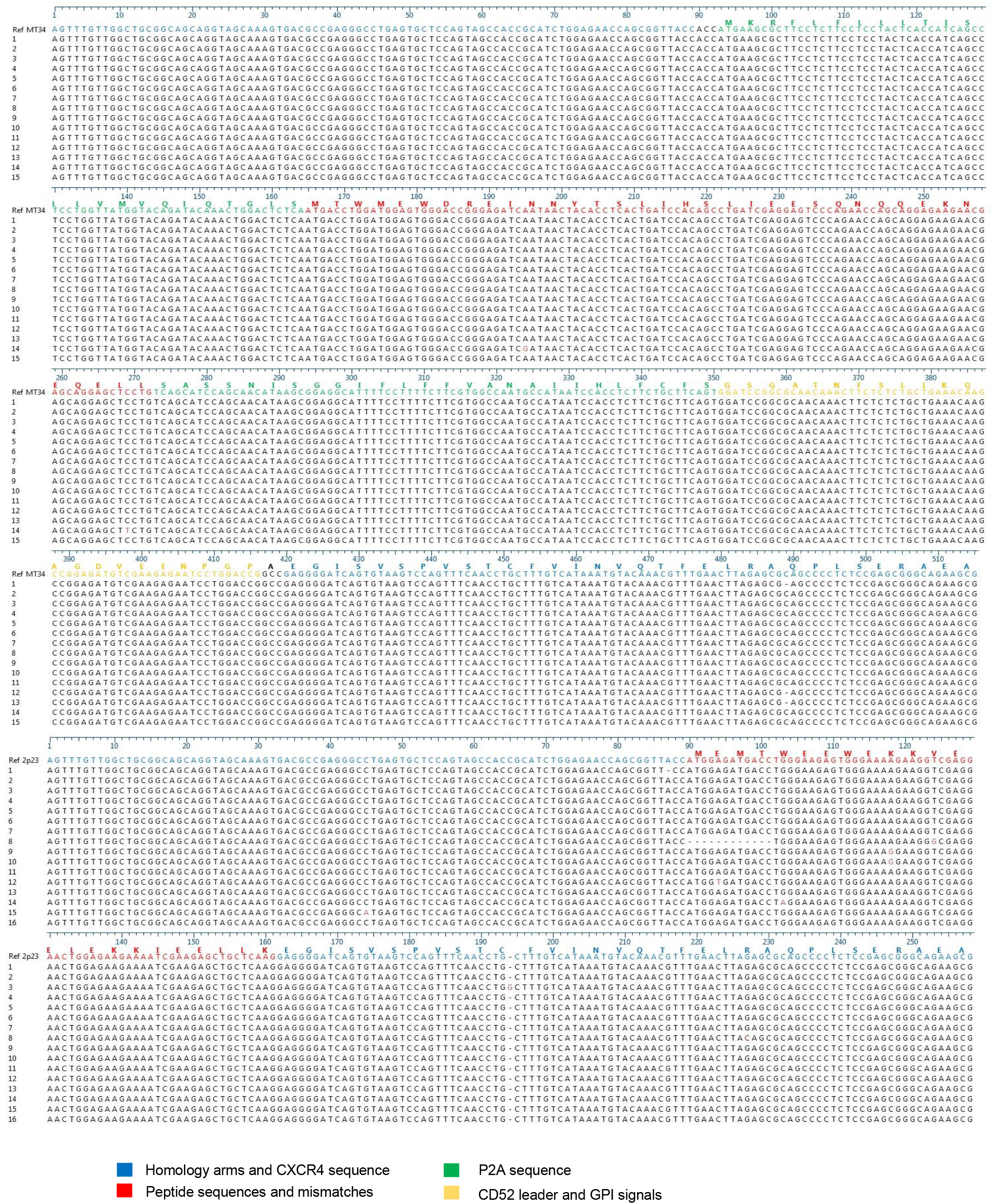
Sequence analysis of CXCR4-integrated DNA products PCR-amplified from MT-C34_GPI_^+^2P23-X4^+^ CEM/R5 cells. PCR-products highlighted by red boxes (Fig.5C) were gel excised, purified, cloned into pJet1.2 vector and analyzed by Sanger sequencing after E.coli transformation and selection of individual bacterial clones.

**Figure S3.**
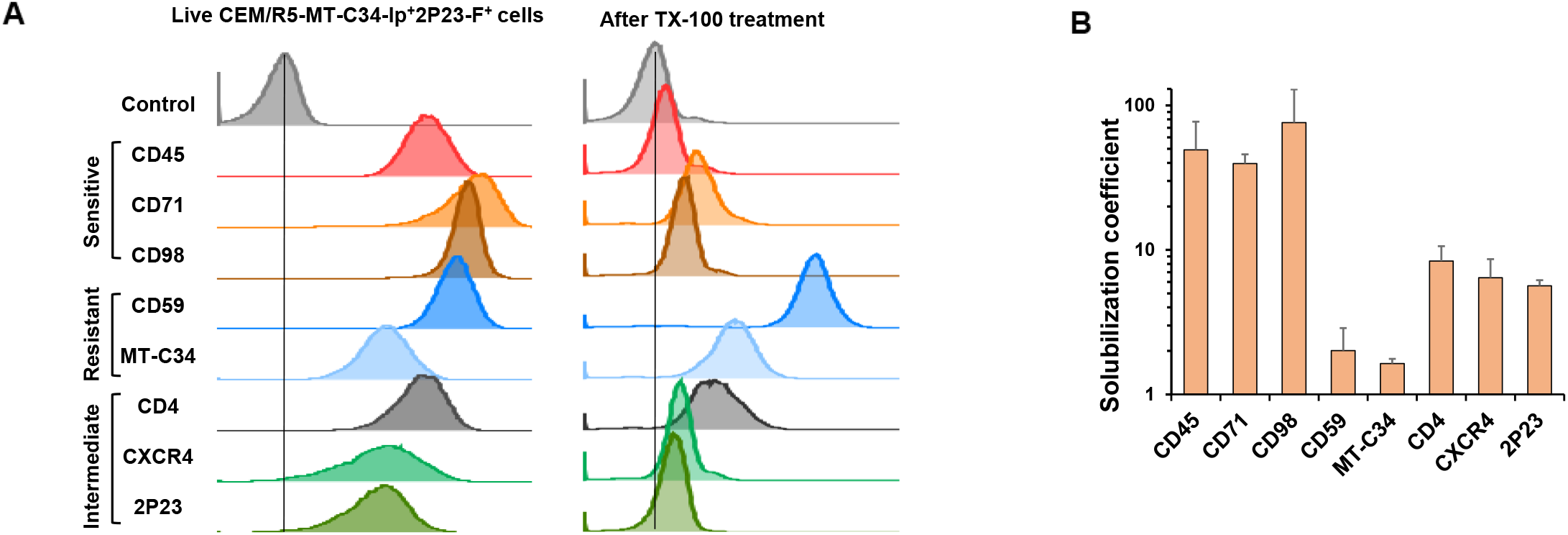
Different sensitivity of lipid raft-associated MT-C34 and 2P23 fused with CXCR4 to Triton X-100 solubilization. **A**. CEM/R5-MT-C34_GPI_^+^2P23-X4^+^ cells were either surface stained with Abs against indicated antigens and the secondary Alexa-488-conjugated Ab (on the left) or treated, first, with TX-100/paraformaldehyde as outlined in Method section, and then the resulting cellular remains were immunostained likewise (on the right). Samples were analyzed by flow cytometry and presented as overlayed histograms with partial off-set. **B**. Mean fluorescence intensities (MFI) measured for each antigen were first adjusted to respective control (without primary Ab). Solubilization coefficients were calculated by dividing adjusted MFIs prior TX-100 treatment to those obtained after TX-100 extraction. Representative histograms and average values with standard deviations obtained from three independent experiments are shown. Non-raft antigens CD45, CD71, and CD98 were easily solubilized by TX-100. The GPI-anchored CD59 and MT-C34 were resistant to detergent treatment, while 2P23 covalently linked to CXCR4 and CD4 displayed intermediate sensitivity to TX-100 solubilization.

**Figure S4.**
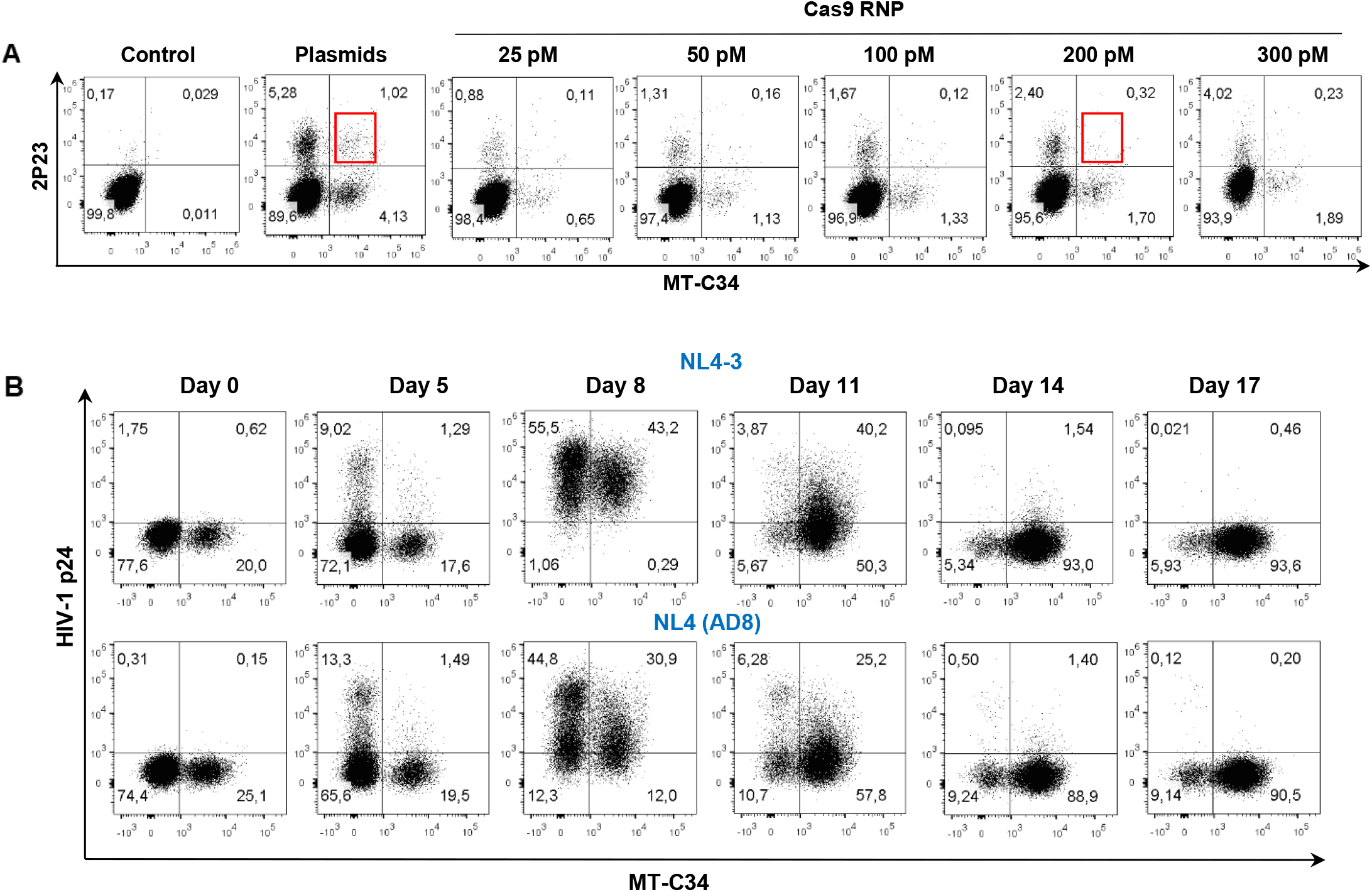
Survival and selective expansion of cells with MT-C34 and 2P23 peptide KI during HIV-1 infection. **A**. Knock-in optimization with Cas9 ribonucleoprotein (RNP) complex. CEM/R5 cells were co-transfected using 1 µg of 2P23-X4, 1 µg of MT-C34_CD52_ PCR-donors, with either gRNA (1 µg) and Cas9 (3 µg) expression plasmids or Cas9 RNP. Double-positive cells (in red boxes) were sorted and used for further analysis. **B**. A culture mixture consisting of 2×10^5^ CEM/R5-MT-C34_GPI_^+^/2P23-X4^+^ cells generated using Cas9 RNP and 8×10^5^ CEM/R5 cells was challenged with 100 ng of X4-tropic NL4-3 or R5-tropic NL4(AD8) strain of HIV-1. The levels of infection spread and the ratios of two cell populations were monitored by flow cytometry after p24 and MT-C34 double-staining. DotPlots from a typical experiment are shown.

**Figure S5.**
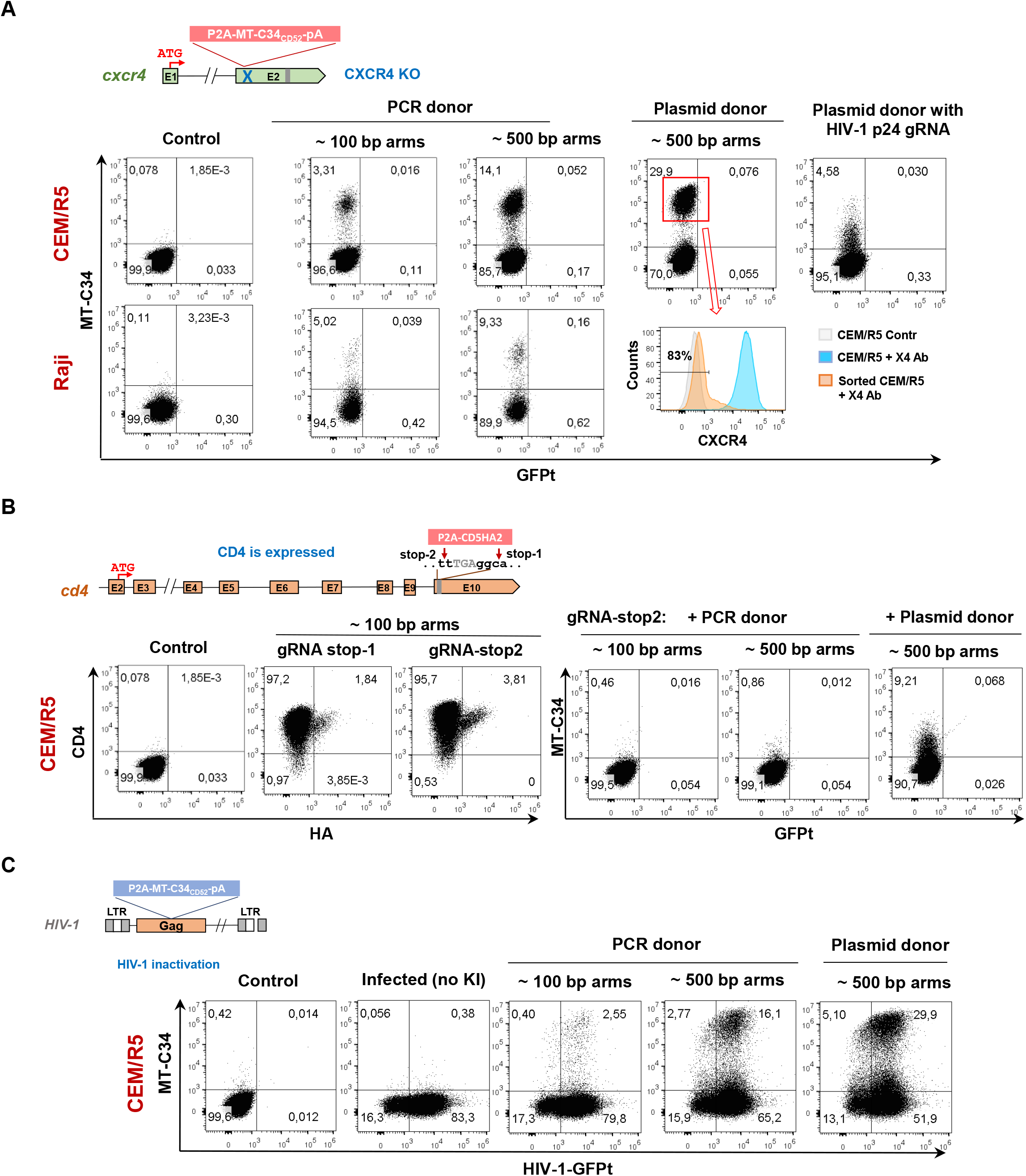
Knock-in efficiency can be improved by homology arm extension, type of DNA donor and gRNA selection. Cells were transfected with 2 pM of PCR or plasmid DNA donor, 1 µg of respective gRNA and 3 µg of Cas9 expression plasmids to initiate KI at loci indicated schematically next to the plotted results (**A** – CXCR4, **B** – CD4 and **C** – HIV-1 integrated DNA). At day 3 posttransfection, cells were immunostained and analyzed by flow cytometry. The levels of targeted gene expression were estimated either at the same time of KI evaluation or followed by positive cell sorting selection and expansion (see histogram). Images are representative of at least two independent experiments.

**Table S1.**
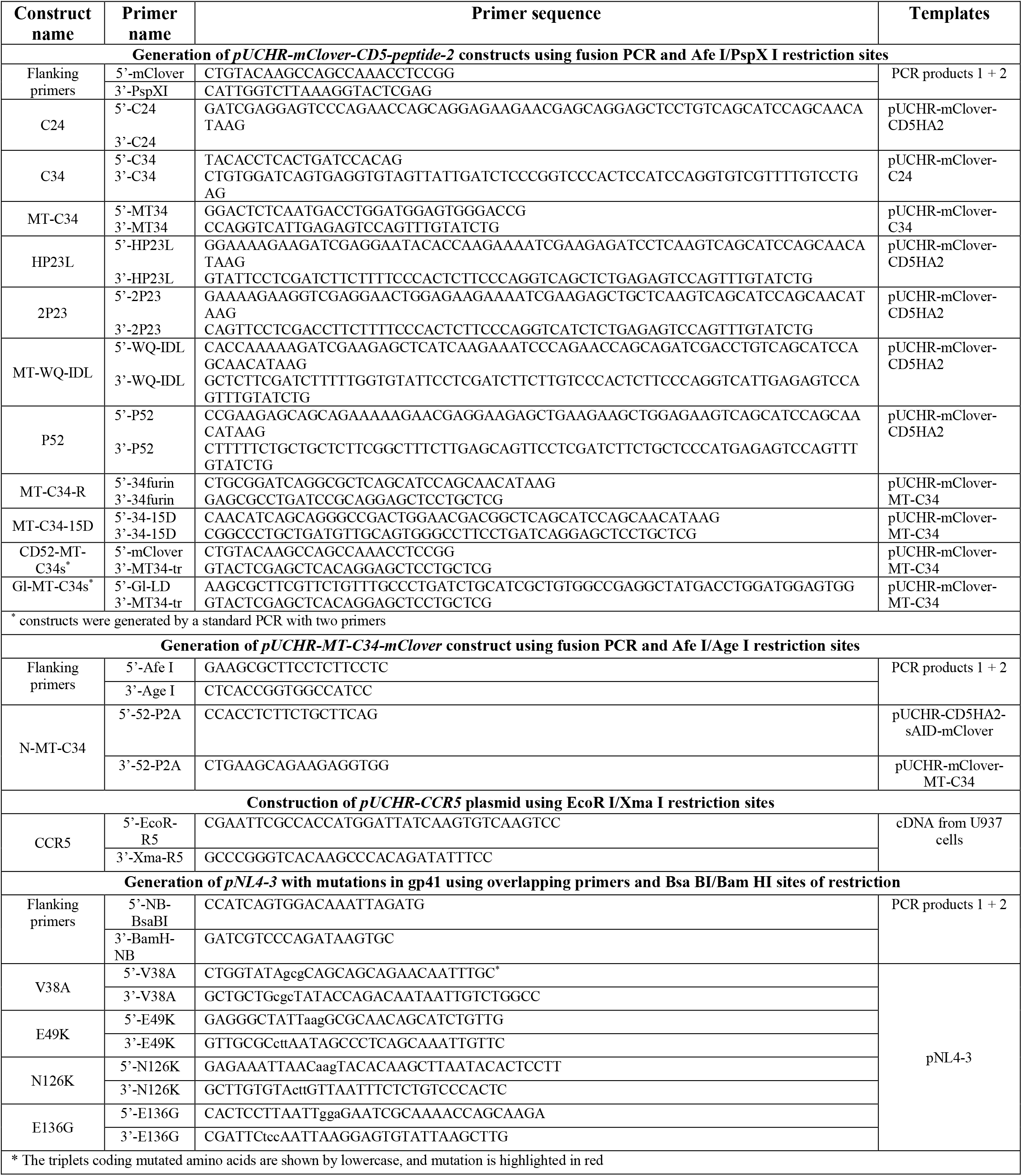
Primers used for plasmid construction.

**Table S2.**
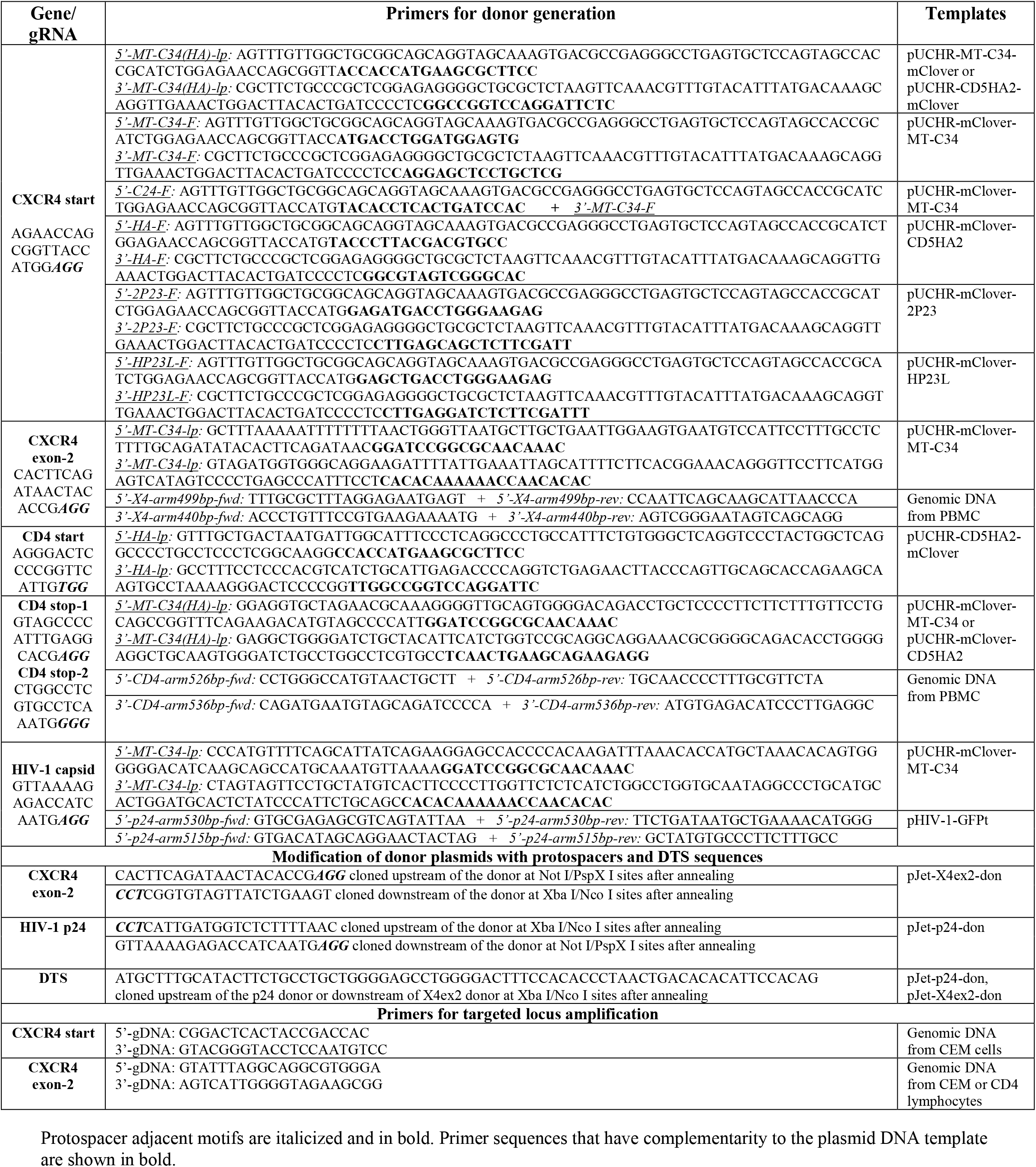
Guide RNA and donor DNA generation.

## Notes

### Competing Interest Statement

The authors have declared no competing interest.

### Summary of Updates

We performed a comparative analysis of the primary CD4 lymphocytes generated using MT-C34 knock-in or lentiviral transduction, created Figure 7 with the new results, and changed the manuscript accordingly. Additionally, we obtained new results on the CXCL12-mediated chemotaxis of CEM cells with KI (Fig.5D). For clarity of study presentation, some construct names were modified, and more descriptions were added to figure legends. Several citations to recent papers were included as well.

